# Neuronal and vascular genome-maintenance states organize opposing multicellular architectures in the aging brain

**DOI:** 10.64898/2026.09.07.749801

**Authors:** Arvin Khoshboresh, Vijay Laxmi Roy, Mofida Abdelmageed, Steffy B. Manjila, Deniz Parmaksiz, Yongsoo Kim, Jesse Gillis, Anirban Paul

## Abstract

Genome maintenance is predominantly studied as a cell-intrinsic process, although its consequences can extend to tissue aging. Brain aging is characterized by region-specific changes in neuronal, immune, and glial cell abundance, altered expression of multiple molecular pathways, and changes in vascular morphology. These features have largely been reported separately, and the molecular relationships underlying regional cellular alterations remain poorly understood. In particular, whether cell-intrinsic genome maintenance is associated with the composition and spatial organization of neighboring cells is unknown. We investigated whether DNA damage response (DDR) transcriptional states provide a molecular link between age-related cellular changes and the surrounding tissue microenvironment. Using MERFISH, we mapped DDR transcriptional programs across ∼600K single cells in matched brain sections from 2–3- and 22-month-old mice. We found that DDR transcriptional states were associated with distinct multicellular neighborhood organizations. Aging also produced region-specific changes in DNA repair pathway module expression and coordination, accompanied by changes in cell-type abundance. Immune enrichment in fiber tracts and oligodendroglial enrichment in the isocortex corroborated established aging patterns, while deep layer 6b near-projecting cortical glutamatergic neurons showed reduced abundance. We developed two complementary spatial analytical approaches to quantify local cell-type composition and distance-dependent neighborhood organization, revealing distinct novel relationships between neuronal and vascular DDR transcriptional states and their surrounding cell populations. Mostly, immune cells were enriched around DDR-low neuronal and vascular cells, “anchors”, while DDR-low neurons additionally showed greater oligodendroglial representation. In contrast, DDR-high vascular anchors were associated with higher proportion of vascular cells proximally and oligodendroglial populations distally, whereas immune cells were enriched at intermediate distances around DDR-low anchors. Together, we identify DDR transcriptional states as an underlying factor likely reshaping multicellular spatial organization, connecting previously separate observations of regional molecular and cellular changes during brain aging. This relationship raises the possibility that altering neuronal or vascular genome maintenance could influence surrounding immune and oligodendroglial neighborhoods. Thus, we establish a novel spatial framework linking cell-intrinsic genome maintenance to the multicellular architecture of the aging brain and provide a foundation for understanding age-related tissue remodeling.

**Graphical Abstract:** 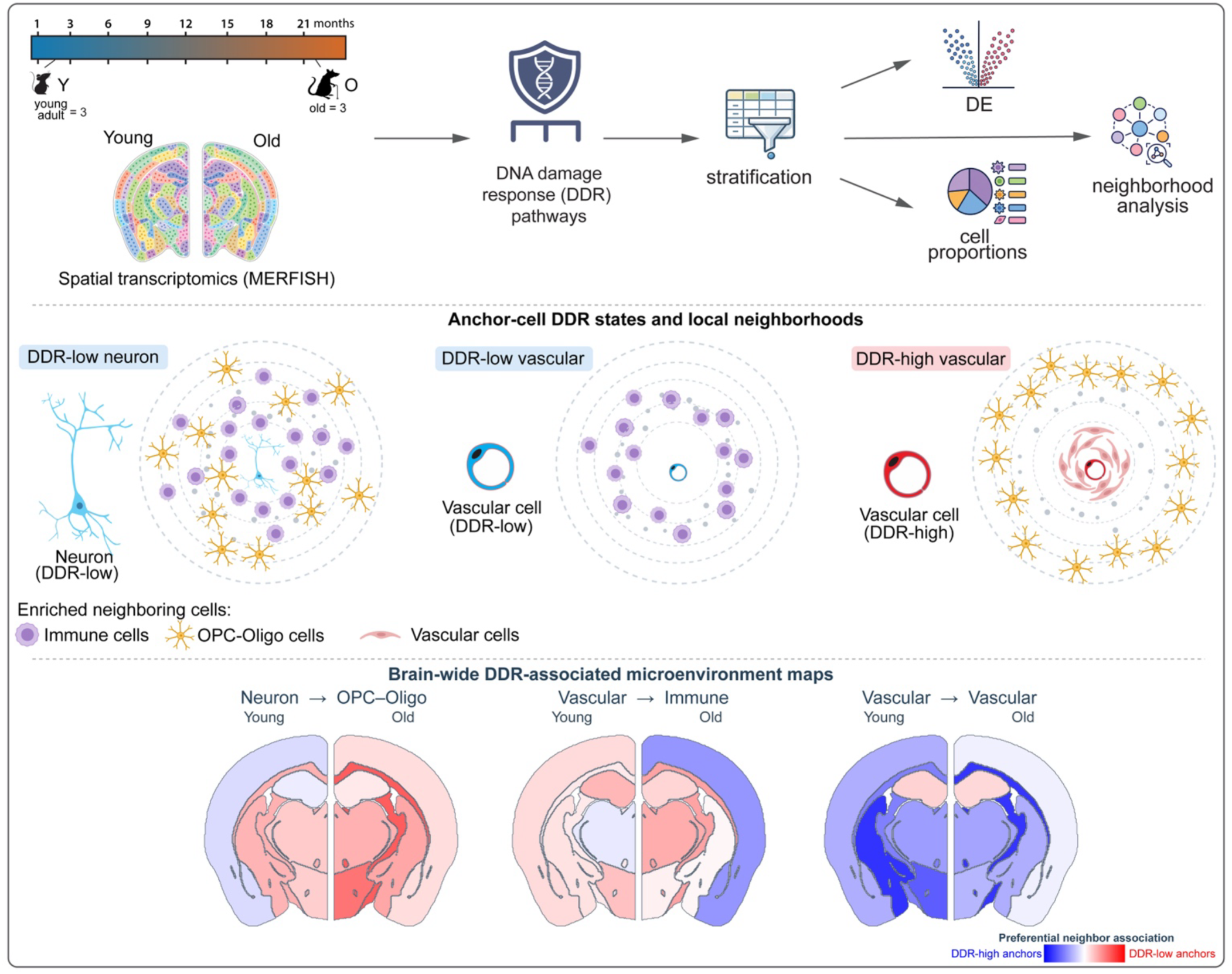

## Introduction

Genome maintenance is a continuous requirement in the adult brain. Long-lived postmitotic neurons sustain oxidative, metabolic and transcription-associated DNA lesions without the dilution afforded by cell division,^1^ while glial, immune and vascular cells respond to distinct proliferative and environmental demands. Persistent damage or maladaptive DNA damage response (DDR) signaling can alter transcription, cellular identity and inflammatory state, linking genome instability to functional decline and neurodegeneration^2,3^. Yet these processes do not unfold in isolated cells. Neurons depend on oligodendroglial metabolism, immune surveillance and the neurovascular unit, each of which changes with age^4–7^.

Transcriptomic studies have established region- and cell-type-specific changes in neuronal, immune and glial populations during brain aging^8–11^. These findings define the cellular landscape in which genome maintenance operates. Connecting that landscape to repair-associated transcription requires measurements that preserve cell identity, anatomical location and the surrounding multicellular environment. This raises a central question: how are regional changes in repair programs related to the cellular organization of aging tissue?

Spatial atlases have established the anatomical organization of brain aging, including age-associated glial and immune states^9^, regional and laminar transcriptional changes, estimated cell-type distributions and inferred interregional ligand–receptor communication^12^. Complementary whole-brain vascular mapping, including light-sheet microscopy, revealed altered vessel geometry and selective reductions in vascular and pericyte density in deep cortical layers, the hippocampal network and basal forebrain^13^. Together, these studies establish regional cellular and vascular remodeling as central features of brain aging. But how these features converge with genome-maintenance states within local tissue environments remains unresolved.

Recurrent cellular neighborhoods provide a framework for addressing this connection. Spatial studies in cancer and intestinal development have established that neighborhood composition and organization capture biological relationships beyond cell abundance^14–17^. In the aging brain, a model linking genome maintenance to tissue remodeling must account for regional repair-program changes, the identity of DDR-expressing cells and the distribution of their neighbors. Defining these relationships would connect molecular maintenance demands to the immune, oligodendroglial and vascular environments in which brain cells age.

Here we used cell-resolved MERFISH spatial transcriptomics^18^ to connect regional DDR expression and coordination with age-associated cellular composition and local neighborhood organization in the mouse brain. Complementary metacell and distance-resolved annular analyses identified genome-maintenance niches as recurrent multicellular environments that surround the individual neuronal or vascular cell, “anchors”, depending on their DDR transcriptional state. These findings provide a link between established cellular and vascular features of brain aging and DNA repair pathway expression state. The resulting model predicts that altering neuronal or vascular repair capacity will remodel distinct immune and oligodendroglial neighborhoods.

## Results

### A spatial framework tests whether genome-maintenance state is associated with tissue organization

Testing a relational model of genome maintenance requires cell identity, anatomy and physical neighborhood to be preserved in the same measurement. Hence we profiled matched anterior and posterior coronal sections from young adult (2-3 months; n = 3) and old (22 months; n = 3) mice by MERFISH on the MERSCOPE platform. Young and old sections were paired within imaging runs, and each animal contributed both sampling levels, yielding 12 spatially resolved sections. This balanced design guarded age comparisons against imaging-run and rostrocaudal sampling effects. A custom panel combined cell-identity markers with genes representing DNA repair and broader genome-maintenance programs (Fig. 1A).

**Figure 1.**
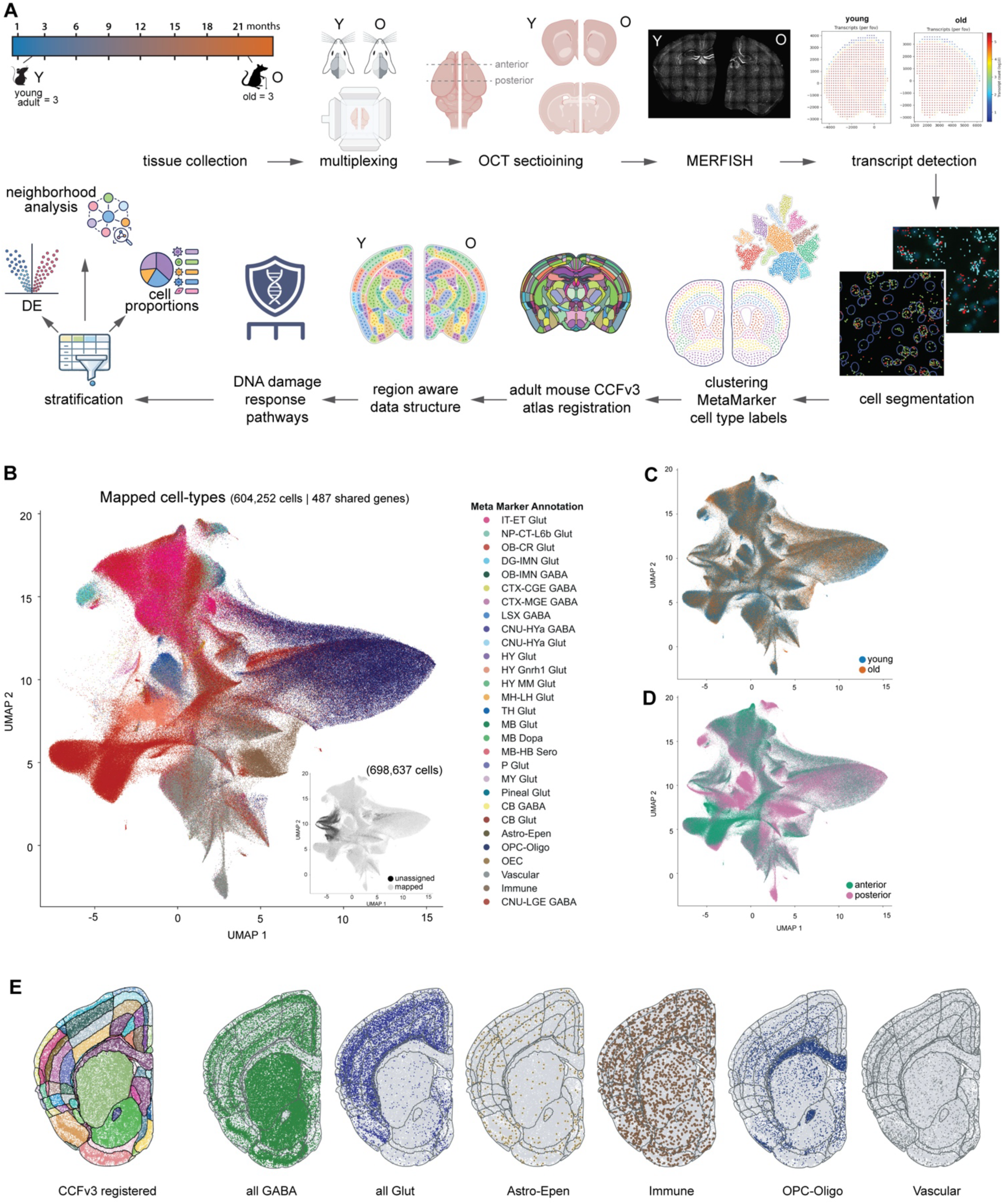
Brain-wide spatial transcriptomic profiling and anatomical registration of young and old mouse brains. A, Experimental and analytical workflow. Coronal brain tissue was collected from young adult (2 months; n = 3) and old (22 months; n = 3) mice, multiplexed to sample anterior and posterior levels, embedded and cryosectioned, and profiled by MERFISH. Detected transcripts were assigned to segmented cells, followed by MetaMarker-based cell-type annotation and registration to the adult mouse CCFv3 anatomical hierarchy. The resulting region-aware, cell-resolved dataset was used to quantify DDR pathway expression and to stratify cells for downstream analyses of differential expression, cell-type composition and spatial neighborhoods. Region- and cell-type-resolved analyses were performed on transcript-count-normalized expression values to distinguish within-cell-type transcriptional changes from changes associated with cellular composition. Anatomical assignments followed the hierarchical organization of the CCFv3 ontology. **B,** UMAP representation of MetaMarker-annotated cells after integration of the spatial datasets. The main embedding shows 604,252 mapped cells across major neuronal and non-neuronal classes using 487 shared genes. Inset, all 698,637 cells before exclusion of cells without a mapped annotation, shown as mapped or unassigned. **C,** UMAP embedding colored by age group, showing the distribution of young and old cells across the integrated transcriptional space. **D,** The same UMAP embedding colored according to anterior or posterior tissue origin, illustrating representation of both anatomical sampling levels across transcriptional populations. E, Representative coronal spatial maps illustrating CCFv3 registration and the distribution of broad cell classes within the registered tissue. From left to right: CCFv3 anatomical assignment, GABA, Glut, Astro-Epen, Immune, OPC-Oligo and Vascular populations. Each point represents a segmented cell positioned at its measured spatial coordinate. Abbreviations: Astro-Epen, astrocyte-ependymal; CCFv3, Allen Mouse Brain Common Coordinate Framework version 3; DDR, DNA damage response; DE, differential expression; GABA, GABAergic; Glut, glutamatergic; MERFISH, multiplexed error-robust fluorescence in situ hybridization; OCT, optimal cutting temperature compound; OPC-Oligo, oligodendrocyte precursor cell-oligodendrocyte; UMAP, uniform manifold approximation and projection; Y, young; O, old.

The initial dataset contained 698,637 segmented cells, of which 604,252 received MetaMarkers-derived identities^19, 20^ were used in subsequent analysis (Fig. 1B). The transcriptional landscape resolved glutamatergic and GABAergic neurons together with astrocyte-ependymal, immune, OPC-oligodendroglial and vascular populations. Young and old cells, as well as anterior and posterior sections, contributed across the principal populations while retaining biological variation associated with age and anatomical origin (Fig. 1C,D).

Registration to the Allen Mouse Brain Common Coordinate Framework (CCFv3)^21^ preserved recognizable neuroanatomy and linked each cell to a hierarchical region while retaining its measured coordinates (Fig. 1E). Each mapped cell was consequently defined by age, animal, transcriptional identity, spatial position and anatomical region. This common coordinate system allowed regional tissue states to be separated from cell-class abundance and enabled direct measurement of the cellular environments surrounding DDR-defined neurons and vascular cells.

### Aging produces region-specific changes in repair-gene expression and coordination

We first mapped age-associated changes in repair-program expression and coordination across anatomical regions. Genome maintenance depends on both the abundance of repair factors and their coordinated deployment. We therefore measured the aging associated change in module expression together with the change in mean within-module Pearson correlation for the overall cellular population of each fine region. Per-gene estimates were pseudobulked at the animal level, age effects were tested by sign-flip permutation of animal-level pseudobulk contrasts and false-discovery rates were controlled across regions. The joint analysis defined coordinated induction (both expression and coexpression are higher in old than young), increased expression with reduced coupling (expression is higher but coexpression is lower in old), concordant suppression and decoupling (both expression and coexpression are lower in old), and coordinated restraint (expression is lower but coexpression is higher in old) (Fig. 2A,C,E,G). Thus, quadrant position represents the direction of age-associated change within each region relative to the young state, not a ranking of absolute expression or coordination across regions.

**Figure 2.**
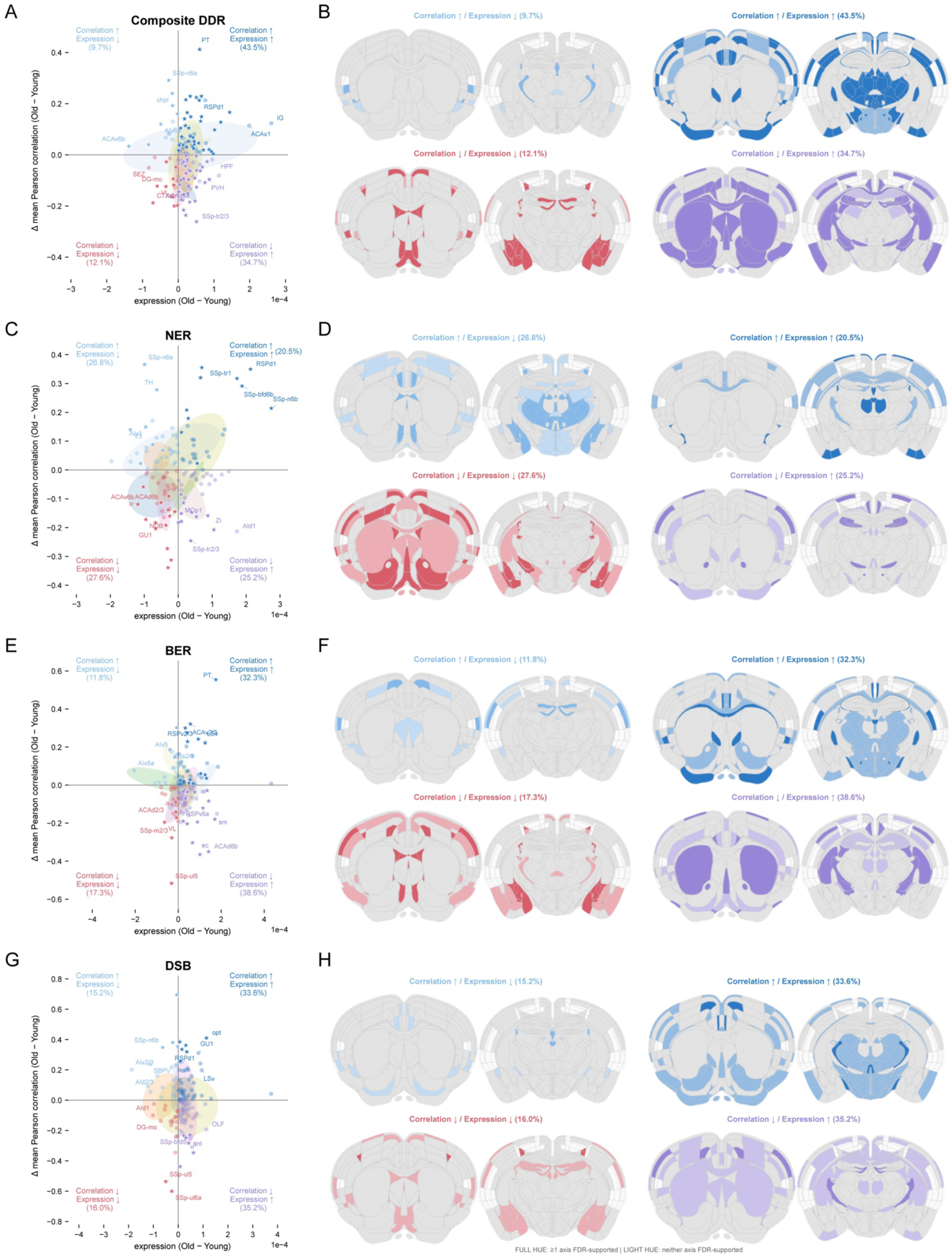
Region-resolved age-associated shifts in DNA damage response pathway expression and coexpression across the mouse brain. A, C, E, **G** DDR module expression-coexpression state space. Each point denotes brain region where all cell labels were included. The x-axis shows the change (old – young) in each module-level expression; the y-axis shows the change (old – young) in mean within-module Pearson correlation. Quadrant colors encode the four signed response states. Quadrant-colored stars denote regions for which expression or coexpression passed the FDR-adjusted threshold (q < 0.05); quadrant-colored circles denote regions for which neither axis passed. A star therefore indicates support for at least one coordinate and does not establish joint significance of the two-dimensional quadrant. Composite DDR is the equal-weight mean of BER, CoreRep, DSB, MMR, NER, and TLS; its support marker indicates that at least one constituent module passed on the relevant axis and is not a composite p-value. Colored ellipses summarize Allen isocortical group structure; percentages report the fraction of all valid regions in each quadrant. Allen brain region “root” was excluded. **B, D, F, H** CCF maps depicting the anatomical regions in the corresponding state-maps in A, C, E, G. For each module, four panels show the anatomical distribution of the same region-level state assignments on Allen CCFv3 slices 460 and 667, rotated clockwise by 90 degrees. Within the displayed quadrant, dark hue shows regions supported on at least one axis by FDR and the same lighter hue denotes regions for which neither axis passed FDR. Territories assigned to the other three quadrants are gray; territories without an assignable region state are white. Percentages report the fraction of valid region assignments in the displayed quadrant. Atlas descendants inherit the assignment of the nearest annotated ancestor for visualization; colored map area is not a statistical weight or region count. **A, B,** Composite DDR. **C, D,** NER. **E, F,** BER. **G, H,** DSB. Abbreviations: BER, base excision repair; CCFv3, Common Coordinate Framework version 3; DDR, DNA damage response; DSB, double-strand break repair; FDR, false discovery rate; NER, nucleotide excision repair.

At the composite DDR level, 97 of 124 regions (78.2%) shifted toward higher expression. Of these, 54 showed coordinated induction and 43 showed increased expression with reduced coupling. Coexpression direction was nearly balanced, increasing in 66 regions and decreasing in 58, and 86 regions received FDR support on at least one axis. Because composite support was assigned when any constituent module passed its threshold, this fraction is a sensitivity indicator rather than a composite statistical test. The dominant tissue-wide pattern was increased DDR expression partitioned between coordinated and less-coupled states, not uniform pathway activation (Fig. 2A,B).

Individual pathways showed distinct regional patterns. NER was distributed comparatively evenly among the four states, whereas BER and DSB were biased toward higher expression (Fig. 2C-H). All FDR-supported DSB regions were supported by coexpression rather than abundance, demonstrating that abundance-only analysis would miss a major dimension of regional DDR organization. The BER bias is consistent with the importance of OGG1-dependent oxidative lesion repair in the aging brain and, more broadly, with coordinated BER regulation and pathway crosstalk, although transcript abundance does not measure repair activity^22–24^. More broadly, the prominence of coexpression changes is consistent with age-dependent remodeling of gene-regulatory coordination^25^.

DDR states were organized at a finer scale than broad anatomical divisions. BER expression and coexpression decreased in dorsal anterior cingulate layers 2/3, whereas NER expression and coexpression increased in dorsal retrosplenial layer 1 (Fig. 2C-F). Adjacent cortical layers frequently occupied different response states, as did neighboring thalamic, hippocampal, cerebral nuclear and hypothalamic territories. Fiber tracts and ventricular interfaces were also prominent: several tracts showed coordinated induction, whereas corpus callosum, external capsule, fimbria and internal capsule showed higher expression with reduced coupling; choroid plexus and third-ventricle territories occupied coordinated-restraint states (Fig. 2B,D,F,H). These patterns extend prior evidence for region-specific repair capacity and the age sensitivity of white matter and periventricular niches^10,11,26^. Because overall-region coexpression can include compositional effects, we interpret this mosaic as a tissue-level substrate ^27^.

### Overall inter-module covariance is preserved across age

Across 123 matched fine regions, we correlated age-specific within-module coexpression values for BER, CoreRep, DSB, MMR, NER and TLS. Young brain contained six FDR-supported module pairs and old brain contained three (Fig. 3A,B), but the overall magnitude of covariance was unchanged (mean signed r, 0.177 young and 0.187 old, P = 0.909; mean absolute r, 0.177 and 0.187, P = 0.899).

**Figure 3.**
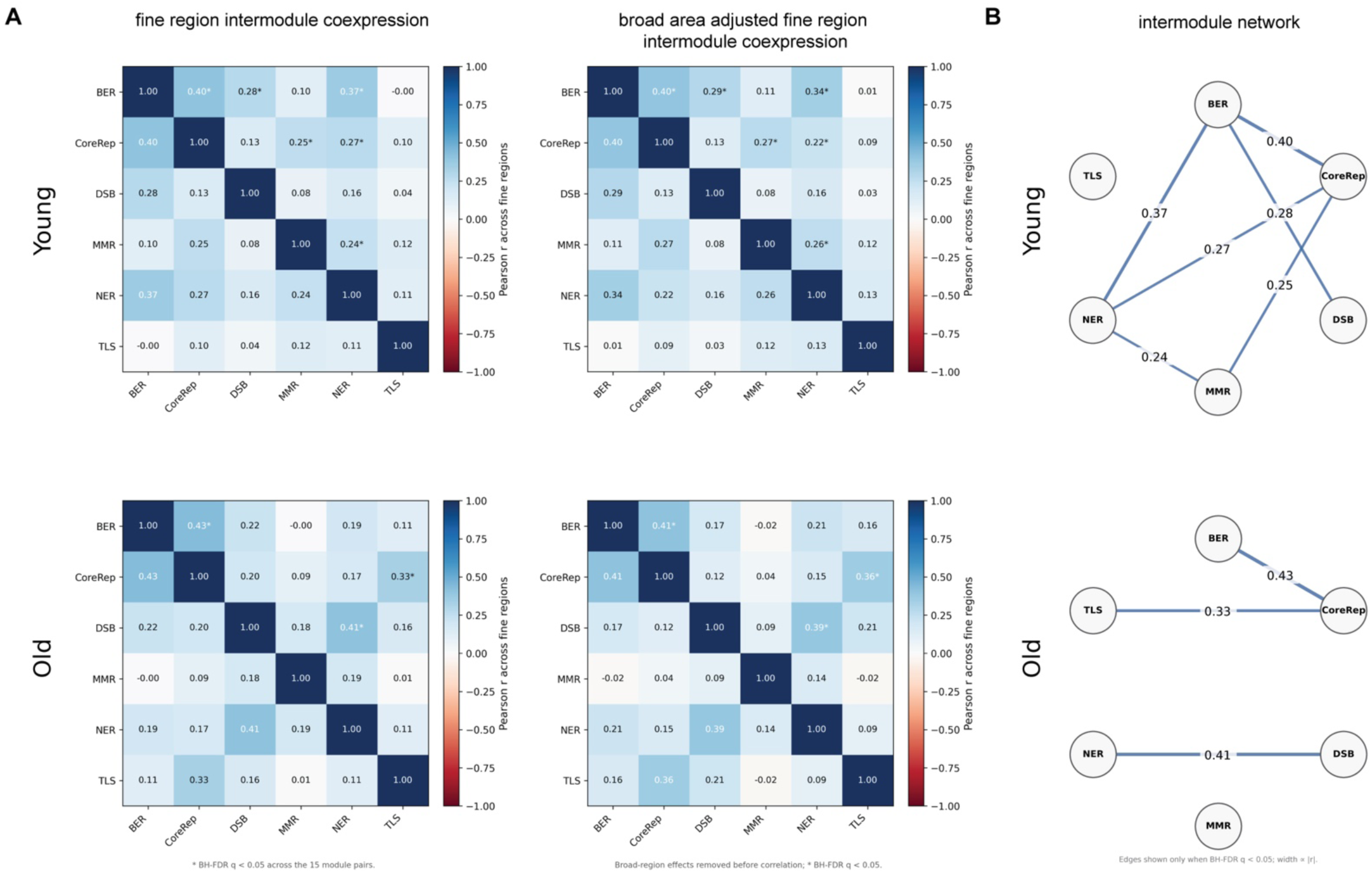
Inter-module DDR covariance is preserved in magnitude across age. A, Heat maps showing pairwise inter-module Pearson correlation across matched fine brain regions in young and old mice. Left, unadjusted fine-region inter-module coexpression. Right, fine-region inter-module coexpression after removal of broad-region effects. Values within cells indicate the correlation coefficient for each module pair. Asterisks denote BH-FDR-significant correlations across the 15 tested unique module pairs. **B,** Network representation of the significant inter-module correlations shown in A. Nodes represent DDR modules and edges indicate BH-FDR-significant positive correlations; edge width is proportional to the magnitude of the correlation coefficient and edge labels show the corresponding r values. These within-age networks describe the FDR-supported covariance structure at each age; direct tests of age differences between individual module pairs are shown in Extended Data Fig. 1. Abbreviations: BER, base excision repair; BH-FDR, Benjamini–Hochberg false discovery rate; CoreRep, core replication/replication-associated program; DDR, DNA damage response; DSB, double-strand break repair; MMR, mismatch repair; NER, nucleotide excision repair; TLS, translesion synthesis.

Individual edges showed candidate redistribution within this preserved magnitude. The largest positive changes were DSB-NER (Δr = +0.254) and CoreRep-TLS (+0.231), and the largest negative changes were BER-NER (−0.174) and CoreRep-MMR (−0.165), but none of the 15 age differences survived FDR correction (Extended Data Fig. 1A). Spearman analysis preserved their directions while reducing the two positive estimates (Extended Data Fig. 1C). Broad-region residualization retained the six young and three old within-age edges, indicating that those covariance patterns were not explained by major anatomical compartments; it did not render any age-difference edge significant (Fig. 3A,B and Extended Data Fig. 1B). Together, these analyses support the preservation of overall covariance magnitude and highlight several directional changes for further investigation.

### Reproducible regional shifts remodel the cellular substrate of DDR states

Regional DDR states are measured in a tissue whose cellular composition also changes with age. We quantified cell-class representation across the mapped brain and within broad anatomical regions (Fig. 4A,B). Proportions were calculated for each sampled section, including zero representation when a sampled region lacked a given cell type, and FDR was controlled across cell types within each region. We further examined consistency across anterior and posterior sections and individual animals so that a regional shift could not be attributed to one sampling plane or one old mouse.

**Figure 4.**
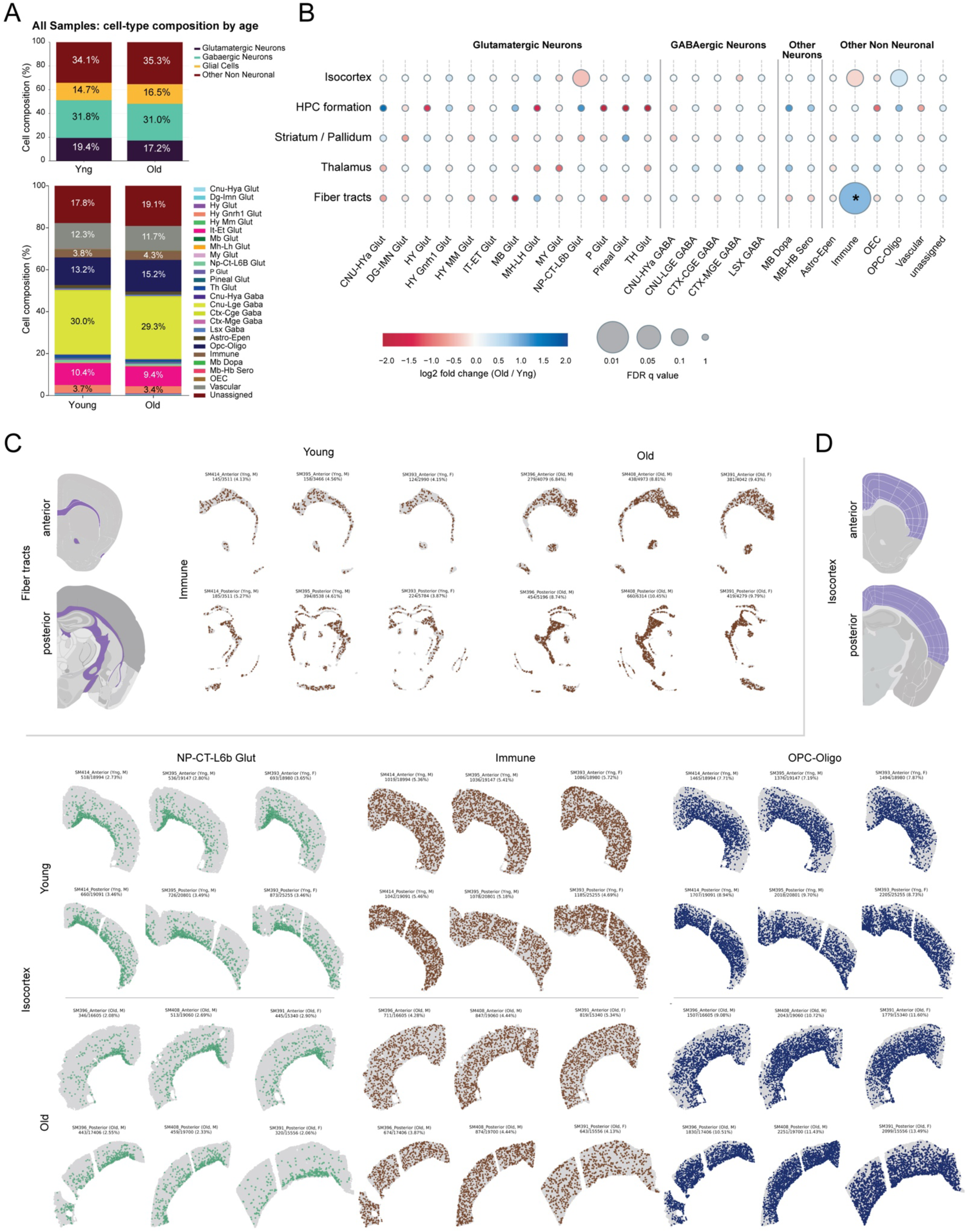
**Age-associated shifts in brain cell-type composition across broad regions**. **A,** Cell-type composition by age across all samples. The upper stacked bars show the aggregate proportions of broad classes in young and old brains, and the lower stacked bars show the corresponding breakdown into finer MetaMarker-derived cell-type labels. Values within bar segments indicate the percentage contribution of each class to the total mapped cell population for each age group. The stacked bars are descriptive pooled-cell summaries and do not define biological n. **B,** Broad-region summary of age-associated compositional change. Rows denote broad anatomical regions and columns denote mapped cell types grouped into glutamatergic neurons, GABAergic neurons, other neurons and other non-neuronal classes. Circle color indicates log2 fold change in relative abundance (Old/Young), and circle size indicates the corresponding FDR q value. This analysis highlights region-specific age-associated enrichment or depletion of individual cell populations, including notable shifts in Fiber tracts and Isocortex. The FDR values summarize section-level comparisons; sections remained linked to their animal of origin and were not treated as additional animals. Panels C and D display the individual sections underlying the highlighted Fiber-tract and Isocortex patterns. C, Individual-section spatial maps for Fiber tracts. Left, CCFv3 masks indicating Fiber tract territory at the anterior and posterior sampling levels. Right, Immune-cell distributions in all sampled young and old sections containing Fiber tract territory, shown separately for anterior and posterior levels. Labels above each map identify the section and report the corresponding section-specific count and percentage. Displaying the complete section-level series allows the age-associated increase in Immune representation within Fiber tracts to be assessed across animals and both anatomical levels. D, Individual-section spatial maps for Isocortex. Top, CCFv3 masks indicating Isocortex territory at the anterior and posterior sampling levels. Bottom, NP-CT-L6b Glut, Immune and OPC-Oligo distributions in all sampled young and old sections, shown separately for anterior and posterior levels. Labels above each map identify the section and report the corresponding section-specific count and percentage. Displaying the complete section-level series allows the regional cellular shifts summarized in B to be assessed across animals and both anatomical levels. Abbreviations: CCFv3, Common Coordinate Framework version 3; FDR, false discovery rate; Glut, glutamatergic; HPC, hippocampal; NP-CT-L6b, near-projecting corticothalamic layer 6b; OEC, olfactory ensheathing cell; OPC-Oligo, oligodendrocyte precursor cell–oligodendrocyte; Yng, young.

Fiber tracts showed a clear immune enrichment that reproduced across anterior and posterior territories and across individual old animals (Fig. 4C). Myelin-rich tracts experience sustained turnover, and age-dependent myelin fragmentation burdens microglial lysosomal clearance and produces white-matter-associated microglial states distinct from those in gray matter^28,29^. The aged isocortex showed a different pattern: OPC-oligodendroglial cells were enriched, whereas NP-CT-L6b glutamatergic and immune populations were depleted (Fig. 4B,D). These opposing immune directions demonstrate regional specialization rather than a uniform brain-wide immune increase^6^. The enrichment of immune cells in fiber tracts and oligodendroglia in cortex is consistent with prior spatial transcriptomic evidence of age-associated increases in white-matter microglia and T cells and in cortical oligodendrocytes^30^.

The isocortical pattern identifies a shifted balance between deep-layer excitatory neurons and the oligodendroglial lineage. Aged OPC states vary by region and niche, and aged microglia can constrain oligodendrocyte generation^5,31^. OPC-oligodendroglial enrichment could reflect altered lineage progression, retention or compensation, while reduced NP-CT-L6b representation could reflect survival or molecular reclassification. These regional shifts establish the cellular context of the repair-program mosaic. Together with published spatial aging observations^30^, our strongest statistically supported changes in cell-type proportions were concentrated in selected anatomical territories rather than spread broadly across the sampled brain. Such localized changes may occur where aging-associated insults, intrinsic molecular alterations, or both are large enough to shift regional cellular composition. However, local neighborhoods may still show differences in composition without large age-associated shifts in cell-type proportions. This motivated a search for recurring DDR-associated microenvironmental patterns in how they change with age between young and old brains. We next tested their local organization by asking how neuronal DDR state relates to immune, oligodendroglial and vascular composition within the same tissue neighborhoods.

### Metacell analysis links low neuronal DDR to immune- and oligodendroglial-rich niches

To test whether neuronal pathway state covaried with the local cellular environment, we next partitioned each section into non-overlapping spatial metacells containing approximately 100 cells. Neuronal transcripts were pseudobulk-averaged within each metacell and used to calculate rank-based DDR scores, whereas immune, OPC-oligodendroglial and astrocyte-ependymal proportions were calculated independently from the cells used for scoring. This separation prevented the abundance of a target cell class from contributing directly to the neuronal DDR metric. Across 50 bins ordered from low to high composite neuronal DDR, OPC-oligodendroglial abundance showed the clearest inverse gradient at both ages. Immune composition showed a smaller within-age gradient but shifted across the DDR-ranked distribution with old age, whereas astrocyte-ependymal differences were modest (Fig. 5A-D). Thus, the metacell analysis most strongly linked low neuronal DDR to oligodendroglial-rich tissue contexts and indicated an age-associated redistribution of immune neighborhoods.

**Figure 5.**
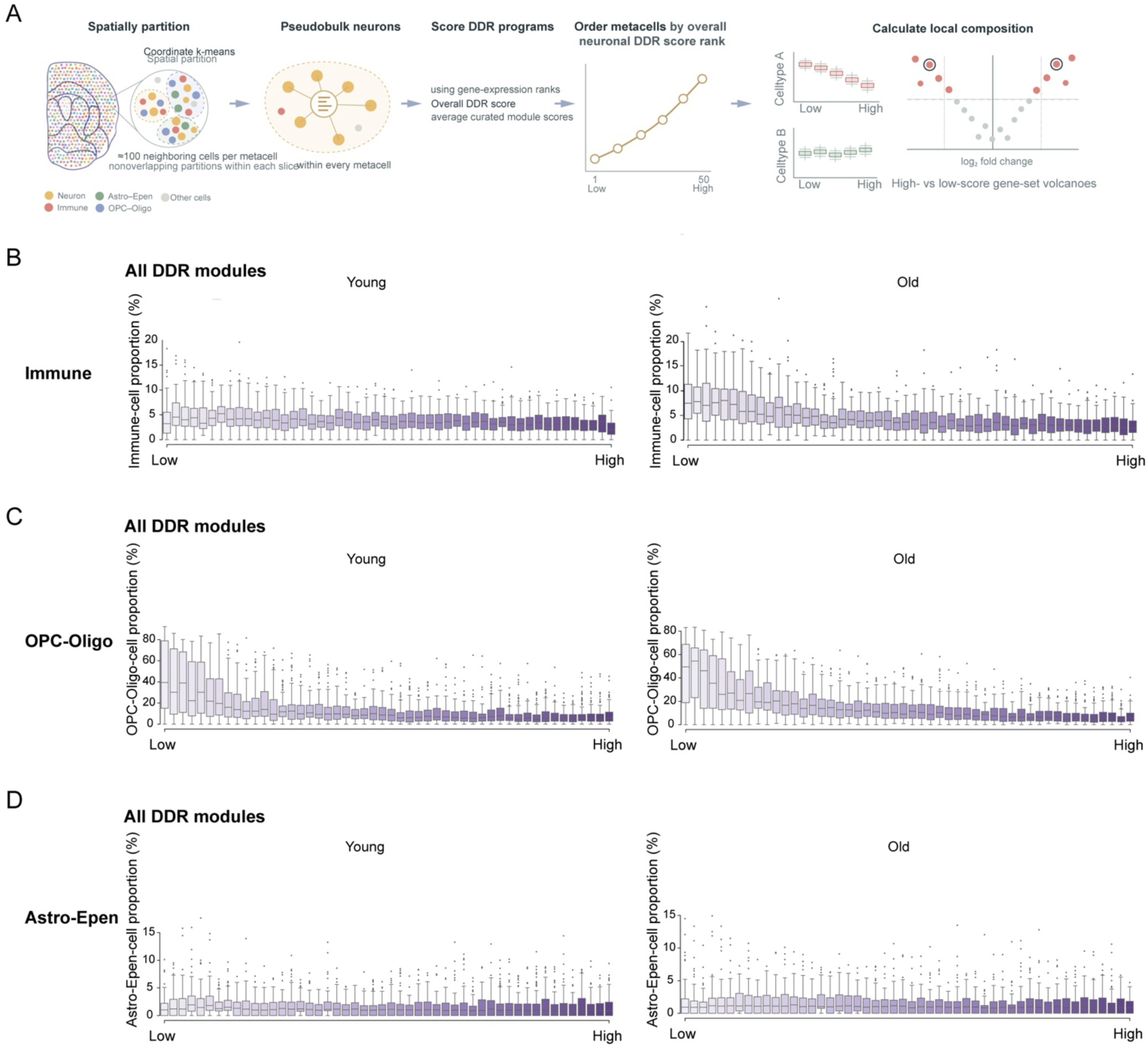
Low neuronal DDR states are associated with distinct non-neuronal spatial microenvironments. A, Schematic of the metacell analysis. Cells within each tissue section were partitioned by spatial proximity into non-overlapping metacells containing approximately 100 neighboring cells. Neuronal expression within each metacell was pseudobulk-averaged and scored using rank-based gene-set scores for the curated DDR programs. An overall neuronal DDR score was calculated from the curated module scores, and metacells were ordered from low to high neuronal DDR score. Local cellular composition was then quantified independently from the pathway-scoring cells. For individual gene sets, metacells were ranked by pathway score, partitioned into 50 bins, and target-cell abundance in the highest- and lowest-scoring bins was compared to generate pathway–composition effect sizes. **B-D,** Distribution of immune (B), OPC-Oligo (C) and Astro-Epen (D) cell proportions across 50 metacell bins ordered from lowest to highest overall neuronal DDR score, shown separately for young (left) and old (right) brains. Box plots show the distribution of target-cell proportions among metacells within each score bin; center lines denote medians and individual outliers are shown. OPC-Oligo showed the clearest inverse gradient with neuronal DDR score in both ages. Immune composition showed a smaller within-age gradient and shifted across the DDR-ranked distribution with age, whereas the Astro-Epen gradient was modest.

Specificity was assessed by extending the analysis to approximately 4,000 Gene Ontology terms, curated DDR modules and descendants of the GO DNA-repair term, with false-discovery control across the complete gene-set universe. In young brain, BER, NER, MMR, DSB and CoreRep showed greater immune representation in low-scoring neuronal metacells; in old brain, all six curated programs showed this direction (Extended Data Fig. 2A). The same orientation extended across broader neuronal programs involving synaptic communication, dendrite organization, xenobiotic response and cell-cycle regulation. We therefore interpret the immune result primarily as an age-associated broadening and redistribution of the neuronal-state relationship, rather than as a large within-age abundance effect.

OPC-oligodendroglial cells showed the strongest and most extensive association. Most DDR modules and numerous DNA-repair descendants were enriched in OPC-oligodendroglial cells within low-scoring neuronal metacells at both ages (Extended Data Fig. 2B). The same direction encompassed calcium and ion transport, neurotransmitter-receptor activity, dense-core vesicle biology, cellular projections, protein-complex assembly and metabolism. Because oligodendroglia support axonal metabolism, this configuration may represent altered axon-glia coupling or increased support requirements^4^. Astrocyte-ependymal abundance was less aligned with composite DDR (Extended Data Fig. 2C). The breadth and consistency of the OPC-oligodendroglial relationship across age and independently tested gene sets established this neuronal gradient as a robust spatial association rather than a feature of any single score.

Metacells thus establish a microenvironmental association at the local level, but they neither identify a focal neuron nor resolve distance. Experimental models show that DSB-bearing or repair-deficient neurons can activate nearby microglia^32,33^ and recently age-associated neuronal states have also been linked to the spatial proximity of nearby microglia^34^, supporting a broader relationship between neuronal and immune-cell organization in the aging brain. So, next we asked whether individual DDR-defined neurons occupied distance-resolved multicellular niches in intact tissue.

### Low-DDR neurons define distance-resolved multicellular niches

We stratified individual neurons at the extremes of each DDR-module score and quantified immune, OPC-oligodendroglial and vascular neighbors across ten annuli spanning 0–150 μm. DDR-low and DDR-high anchors were defined by the 6th and 94th percentiles, and each annulus was tested with multiplicity correction before integration into an age-specific radial association score. As a compositional control, broad cell-class representation within DDR-low and DDR-high pools was observed to be similar across young and old brains and across individual modules and composite DDR (Extended Data Fig. 3), arguing against a gross imbalance in neuronal, glial, immune or vascular representation as an explanation for the annular patterns. Positive radial association scores indicate enrichment around DDR-low neurons and negative scores indicate enrichment around DDR-high neurons. The old-minus-young difference between these age-specific scores is the age-polarization score (APS); we used APS only to summarize displacement between profiles, and based the biological conclusions on the age-specific directions and corrected annulus-level comparisons. Supplementary Fig. 6 provides a schematic guide to interpreting these annular contrasts and age-specific radial scores.

Immune neighborhoods were regionally polarized around neuronal DDR state (Fig. 6 and Extended Data Fig. 4). Thalamus and hypothalamus generally shifted toward stronger immune association with DDR-low neurons, whereas cortex moved toward DDR-high neuronal states. Hippocampal immune neighborhoods remained predominantly DDR-low-associated, with smaller and module-dependent age effects. Representative profiles separated an overall age-related increase in local immune fraction from preferential partitioning of that environment around DDR-low neurons (Fig. 6A-C). Thus, density and DDR orientation were distinct properties of the same neighborhood.

**Figure 6.**
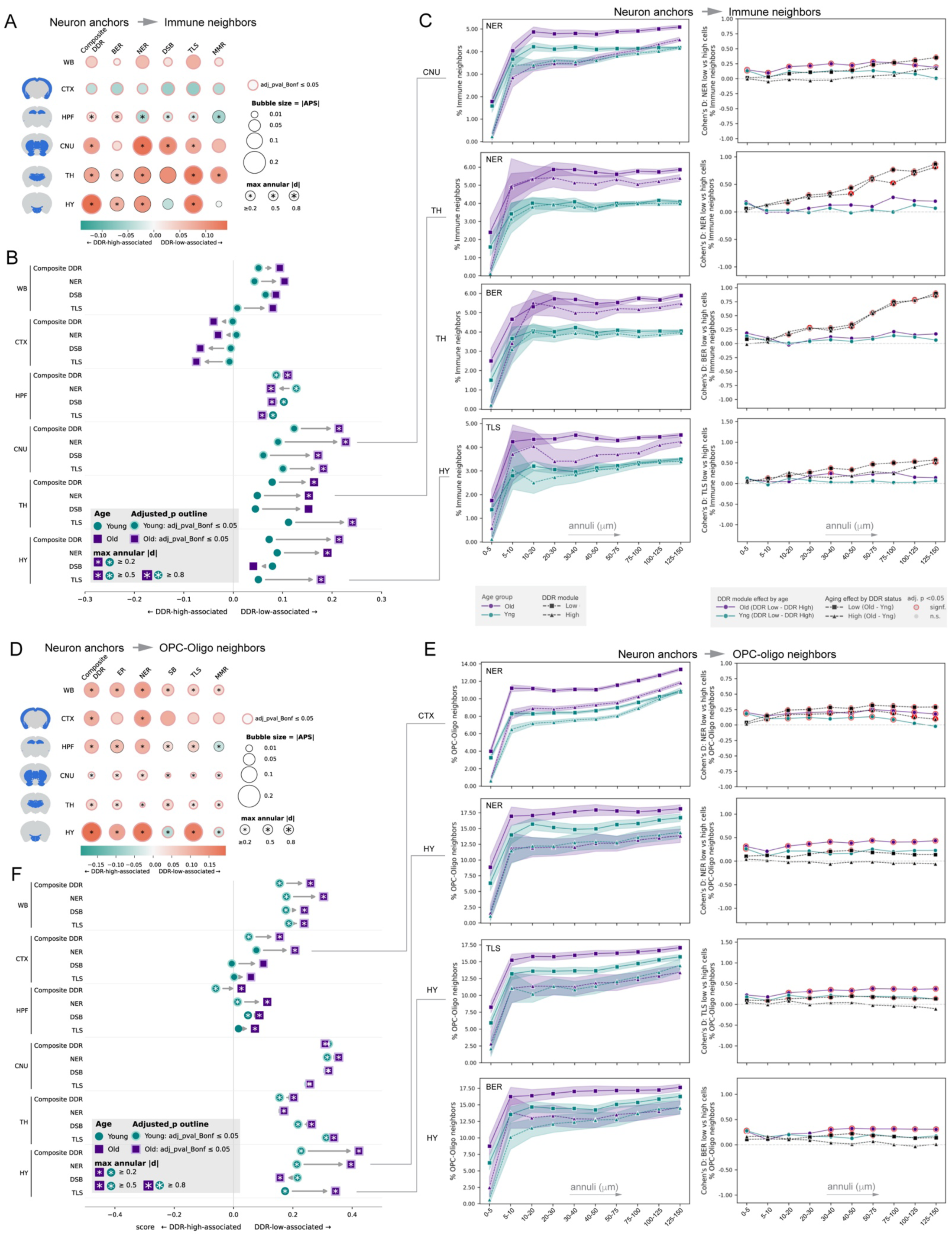
Aging reorganizes neuron-centered immune and oligodendroglial neighborhoods by neuronal DNA damage response state A, Brain-wide summary of age-dependent polarization of immune-cell neighborhoods surrounding neuronal anchors. For each brain region and DDR module, annulus-level Cohen’s d values comparing neighbor abundance around DDR-low versus DDR-high neuronal anchors were integrated across the 0–150 μm radial neighborhood to yield an age-specific radial association score. The age-polarization score (APS) was calculated as the old radial association score minus the young radial association score. Positive APS values indicate an age-associated shift toward DDR-low-associated neighborhoods, whereas negative values indicate a shift toward DDR-high-associated neighborhoods. Bubble area is proportional to |APS|. Bubble outlines denote Bonferroni-adjusted significance in the underlying annulus-level analysis, and stars denote the maximum absolute annulus-level Cohen’s d (|d| ≥ 0.2, ≥ 0.5 or ≥ 0.8). Columns show Composite DDR, BER, NER, DSB, TLS and MMR across WB, CTX, HPF, CNU, TH and HY. **B,** Directional young-to-old changes in neuron-to-immune radial association for Composite DDR, NER, DSB and TLS. Teal circles and purple squares indicate young and old age-specific radial association scores, respectively, with arrows indicating the young-to-old transition. Negative values indicate preferential immune association with DDR-high neuronal anchors and positive values indicate preferential association with DDR-low neuronal anchors; trajectories crossing zero therefore indicate an age-dependent reversal of DDR-state association. Marker outlines denote Bonferroni-adjusted significance in at least one underlying annulus, and stars indicate the maximum absolute annulus-level Cohen’s d using the thresholds in the grey inset. **C,** Representative distance-resolved neuron-to-immune relationships illustrating regional and pathway-specific patterns identified in **A, B**: CNU–NER, TH–NER, TH–BER and HY–TLS. Left, percentage of immune neighbors surrounding DDR-low and DDR-high neuronal anchors in young and old mice across successive non-overlapping annuli from 0–5 to 125–150 µm; shaded regions indicate 95% confidence intervals. Right, corresponding annulus-level effect-size profiles. Colored lines show Cohen’s *d* for DDR-low versus DDR-high neuronal anchors separately in old and young animals, with positive values indicating DDR-low-associated and negative values DDR-high-associated immune neighborhoods. Black dashed lines show old-versus-young Cohen’s *d* separately within DDR-low and DDR-high neuronal anchors. Red-outlined points indicate Bonferroni-adjusted *P* < 0.05; gray points are not significant. **D,** Brain-wide summary of age-dependent polarization of OPC-oligodendroglial neighborhoods surrounding neuronal anchors. APS, bubble size, color, significance outlines and maximum annulus-level effect-size symbols are encoded as in A. Positive values indicate an age-associated shift toward preferential OPC-oligodendroglial association with DDR-low neuronal anchors and negative values indicate a shift toward DDR-high neuronal anchors. **E,** Representative distance-resolved neuron-to-OPC-oligodendroglial relationships: CTX-NER, HY-NER, HY-TLS and HY-BER. Left, percentage of OPC-oligodendroglial neighbors surrounding DDR-low and DDR-high neuronal anchors across the 0–150 μm annuli, stratified by age. Right, corresponding DDR-low versus DDR-high Cohen’s d profiles and old-versus-young effect-size profiles, displayed and annotated as in C. These plots resolve whether DDR-state and aging effects are proximal, spatially persistent or vary with radial distance. **F,** Directional young-to-old changes in neuron-to-OPC-oligodendroglial radial association for Composite DDR, NER, DSB and TLS. Young and old age-specific radial association scores are displayed as in B. Arrow direction and length denote the direction and magnitude of age-associated spatial reorganization, respectively; rightward shifts indicate increasing association with DDR-low neuronal neighborhoods and leftward shifts indicate increasing association with DDR-high neuronal neighborhoods. The bubble and directional plots summarize the underlying annulus-level effect-size profiles; statistical annotations derive from the corresponding distance-resolved comparisons. Age-specific radial association scores and APS were calculated from the unfiltered annulus-level Cohen’s d profiles as described above. Young mice were 2 months old and old mice were 22 months old; the analysis comprised anterior and posterior coronal sections from three animals per age group (six sections per age group). Abbreviations: APS, age-polarization score; BER, base excision repair; CNU, cerebral nuclei; CTX, cerebral cortex; DDR, DNA damage response; DSB, double-strand break repair; HPF, hippocampal formation; HY, hypothalamus; MMR, mismatch repair; NER, nucleotide excision repair; OPC, oligodendrocyte precursor cell; TH, thalamus; TLS, translesion synthesis; WB, whole brain.

OPC-oligodendroglial neighborhoods showed a more consistent pattern. Associations with DDR-low neurons were present in young brain and generally strengthened with age in whole brain, cortex, thalamus and hypothalamus. Cerebral nuclei retained strong relationships at both ages, producing modest APS because aging preserved rather than created the association. The hippocampal composite crossed from a DDR-high bias in young brain to a weak DDR-low bias in old brain (Fig. 6D-F). Vascular neighbors followed a third, regionally conditional pattern: whole-brain and cortical relationships shifted toward DDR-low neuronal states, hippocampal formation and cerebral nuclei generally preserved DDR-low associations, and thalamus remained DDR-high-associated (Extended Data Fig. 4A,B).

The annular analysis recovered the same association observed using complementary spatial metacell analysis and added distance and anatomical resolution. Neuronal DDR state was associated with the identity and radial distribution of nearby non-neuronal cells. The association of immune and OPC-oligodendroglial cells with DDR-low neurons therefore reflected their distribution around defined neuronal anchors rather than a uniform increase in non-neuronal density. We next extended this framework to vascular anchors to connect DDR-associated neighborhoods with the neurovascular environment.

### Vascular DDR states reveal region-specific and radially ordered neighborhoods

To separate pathway-state effects from anchor-cell identity, we compared DDR-associated spatial organization across neuronal and vascular anchors. As a developmentally and functionally distinct comparator to the neural parenchyma, we examined the vascular compartment, whose endothelial lineage arises outside the neuroectoderm and assembles with mural and perivascular cells during CNS vascular development^35,36^. Vascular cells coordinate perfusion, barrier integrity and cellular exchange across the neurovascular unit^7,13,37^. We repeated the annuli analysis with vascular cells as anchors using the same scoring thresholds, annular geometry, effect-size orientation and multiplicity correction (Fig. 7). This matched analysis allowed the neuronal and vascular patterns to be compared using the same analysis parameters.

**Figure 7.**
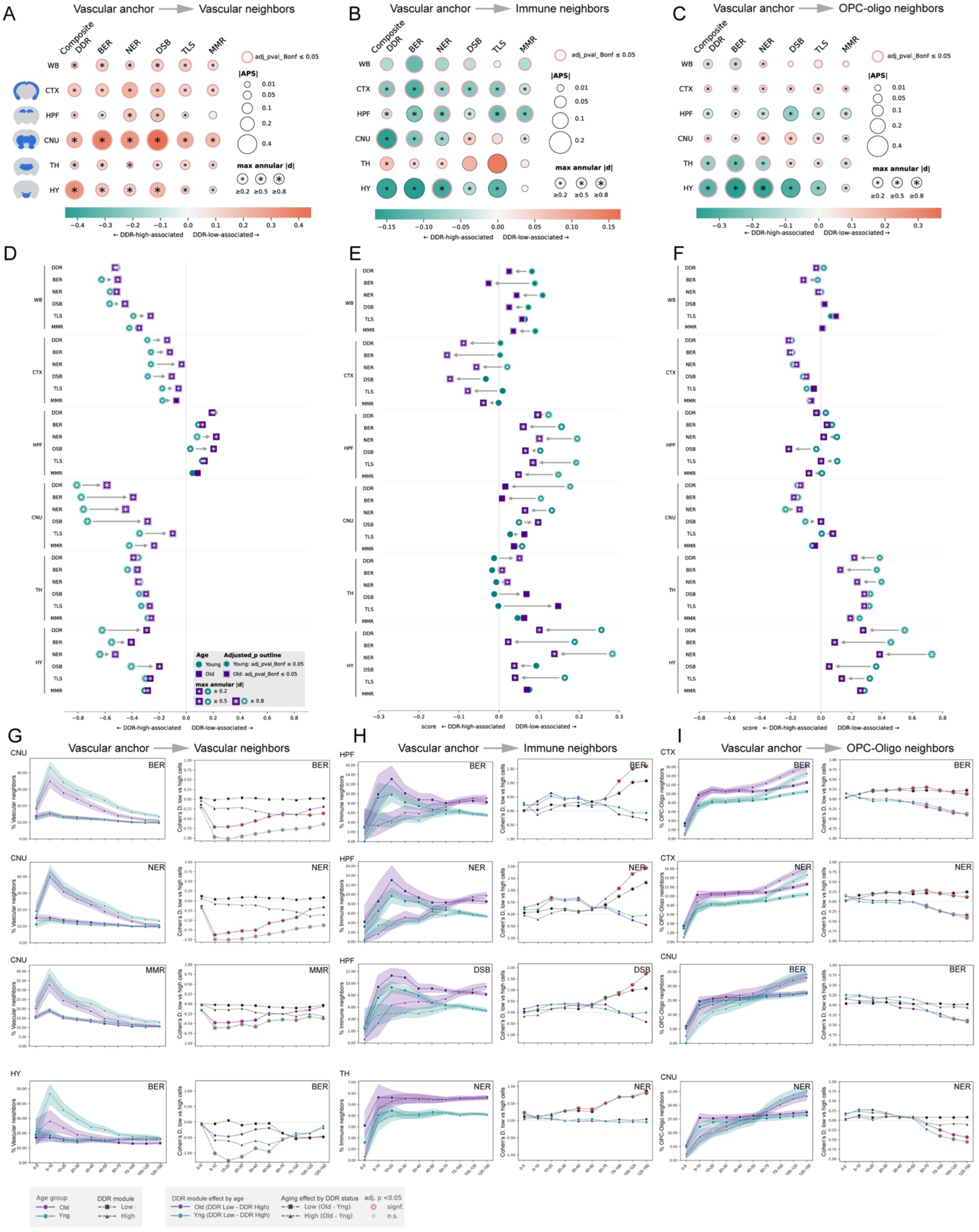
Aging reorganizes vascular-centered cellular neighborhoods according to vascular DDR state. A-C, Bubble-plot summaries of age-associated changes in vascular anchor-to-vascular neighbor (A), vascular anchor-to-immune neighbor (B) and vascular anchor-to-OPC-oligodendroglial neighbor (C) relationships across WB, CTX, HPF, CNU, TH and HY. Vascular anchors were stratified by the indicated module score, and neighbor proportions were quantified in ten non-overlapping annuli spanning 0–150 μm. For each age, all finite annulus-level Cohen’s d values for DDR-low versus DDR-high anchors were multiplied by the corresponding annulus width, summed and divided by the total sampled radial width to yield the age-specific radial association score; no effect-size or significance filtering was applied. Negative scores indicate greater neighbor representation around DDR-high vascular anchors, whereas positive scores indicate greater representation around DDR-low vascular anchors. The age-polarization score (APS) was calculated as the old radial association score minus the young radial association score. Bubble color shows signed APS, with teal indicating an age-associated shift toward DDR-high vascular anchors and salmon indicating a shift toward DDR-low vascular anchors; bubble area is proportional to |APS|. Color and size scales are shown separately for each relationship. A red outline indicates that at least one contributing annulus in either age group had a Bonferroni-adjusted two-sided Mann-Whitney U-test P < 0.05. Black star size denotes the maximum absolute annulus-level Cohen’s d across the contributing age-specific comparisons: small, |d| ≥ 0.2; medium, |d| ≥ 0.5; and large, |d| ≥ 0.8. **D-F,** Directional dumbbell plots showing the young-to-old change in the age-specific radial association score for vascular-to-vascular (D), vascular-to-immune (E) and vascular-to-OPC-oligodendroglial (F) relationships. Arrows visualize direction of chronological aging and do not imply the direction of signaling. Teal circles denote young and purple squares denote old, and grey arrows connect the paired values in the young-to-old direction. Rightward arrows indicate an age-associated shift toward DDR-low vascular anchors, leftward arrows indicate a shift toward DDR-high vascular anchors and trajectories crossing zero indicate reversal of the predominant DDR-state association. Light teal and light purple marker outlines identify young and old scores, respectively, for which at least one annulus had Bonferroni-adjusted P < 0.05. White star size denotes the maximum absolute annulus-level Cohen’s d within the corresponding age group using the thresholds defined above. These outlines and stars annotate the contributing annulus-level comparisons and do not represent an independent significance test of APS or of the young-to-old difference. **G-I,** Representative distance-resolved profiles underlying the integrated relationships. G, Vascular-to-vascular profiles for CNU BER, NER and MMR and HY BER. H, Vascular-to-immune profiles for HPF BER, NER and DSB and TH NER. I, Vascular-to-OPC-oligodendroglial profiles for CTX BER and NER and CNU BER and NER. In the left plot of each pair, lines show the percentage of the indicated neighbors across successive annuli for DDR-low and DDR-high vascular anchors in young and old animals; shaded bands denote 95% confidence intervals. Purple and teal indicate old and young, respectively, whereas squares and triangles denote DDR-low and DDR-high anchor states. In the corresponding right plots, colored lines show annulus-specific Cohen’s d for DDR-low versus DDR-high anchors within each age group, and black lines show the old-minus-young effect within the DDR-low and DDR-high anchor states. Red-outlined points indicate Bonferroni-adjusted two-sided Mann-Whitney U-test P < 0.05 for the corresponding annulus-level contrast; grey points are not significant. Arrows in the panel headings indicate analytical conditioning from anchor to neighbor and do not imply signaling direction. Young mice were 2 months old and old mice were 22 months old; the analysis comprised anterior and posterior coronal sections from three animals per age group (six sections per age group). **Abbreviations:** APS, age-polarization score; BER, base excision repair; CI, confidence interval; CNU, cerebral nuclei; CTX, cerebral cortex; DDR, DNA damage response; DSB, double-strand break repair; HPF, hippocampal formation; HY, hypothalamus; MMR, mismatch repair; NER, nucleotide excision repair; n.s., not significant; OPC-Oligo, oligodendrocyte precursor cell–oligodendrocyte group; TH, thalamus; TLS, translesion synthesis; WB, whole brain.

Immune enrichment around DDR-low anchors was shared across neuronal and vascular compartments. OPC-oligodendroglial enrichment showed a different relationship: it predominantly favored DDR-low neurons but favored DDR-high vascular anchors in cortex and cerebral nuclei. Vascular neighbors also favored DDR-high vascular anchors (Fig. 7A-C). Together, these findings link neighborhood composition to the combination of anchor-cell identity and DDR transcriptional state.

Distance-resolved analysis revealed radial organization around vascular anchors. Vascular neighbors formed the most proximal and consistently DDR-high-associated component across most modules in whole brain, cortex, cerebral nuclei, thalamus and hypothalamus, with hippocampal formation as a notable exception. Aging generally shifted this relationship toward DDR-low anchors, but more often attenuated than reversed the underlying DDR-high association (Fig. 7A,D,G). Immune neighborhoods showed the opposite pattern across whole brain, hippocampal formation, cerebral nuclei and hypothalamus, although the association weakened with age in several contexts. Cortex shifted toward DDR-high-associated immune neighborhoods, whereas thalamus moved toward DDR-low anchors (Fig. 7B,E,H).

OPC-oligodendroglial organization separated the neural parenchyma from the neurovascular interface most clearly. Cortex and cerebral nuclei favored DDR-high vascular anchors, matching the vascular-neighbor direction and opposing the prevailing immune bias. Hippocampal formation, thalamus and hypothalamus instead favored DDR-low anchors. Representative cortical and cerebral-nuclear profiles separated across proximal, intermediate and distal annuli (Fig. 7C,F,I). Vascular DDR state was associated with proximal vascular neighbors, immune cells at intermediate range and more distal, region-specific oligodendroglial neighbors.

These results establish anchor identity and neighbor distance as dimensions of DDR-associated vascular organization. DDR-high and DDR-low describe relative transcriptional states; damage burden and repair capacity are separate functional measurements. Vascular aging includes endothelial senescence^38^, altered DDR signaling^39^ and impaired pericyte remodeling^40^, while endothelial repair deficiency disrupts barrier function^41^. The observed neighborhoods connect vascular DDR transcriptional states to the local cellular environments in which these processes occur.

## Discussion

Regional changes in repair-gene expression and coordination occurred alongside shifts in neuronal, immune and oligodendroglial representation. At the local scale, DDR transcriptional states were associated with reproducible neighborhood composition and distance-dependent organization around neuronal and vascular anchors that changed between young and old brains. We propose that changes in neuronal and vascular repair capacity may differentially influence inflammatory signaling and trophic support, linking genome stress to the cellular remodeling of aging tissue. The observed broader regional variations in DNA repair module expression and coordination with aging, regional cell composition differences, and the altered microenvironment organization are manifestation of a common framework.

BER expression and coexpression decreased in dorsal anterior cingulate layers 2/3, whereas both measures increased for NER in dorsal retrosplenial layer 1. These contrasting responses place repair-pathway regulation within specific cortical territories rather than a uniform brain-wide trajectory. Alongside preserved overall inter-module covariance, they identify regional remodeling of expression and coordination within a broadly maintained relationship among repair programs. Prior evidence for region-specific repair capacity and the role of oxidative lesion repair in brain aging supplies a rationale for investigating these spatial differences^22,26^. The model predicts that the consequences of local genome stress depend jointly on the affected repair program and the surrounding cellular environment.

The fine-scale cellular findings then connect our observations to recent spatial and human aging reports. Immune enrichment in fiber tracts and oligodendrocyte enrichment in isocortex parallel the age-associated increases in T cells and microglia in white matter tracts and oligodendrocytes in cortex reported by Sun et al. (2025). Their spatial aging clocks also associated T-cell proximity with accelerated transcriptomic aging in neighboring cells^30^. These findings, together with immune enrichment around DDR-low anchors here, suggest a potential relationship between inflammatory microenvironments and local genome-maintenance states during aging. Yang et al. (2026) reported late-life glial immune programs in the human prefrontal cortex, with DNA repair emerging among several pathways enriched in convergent neuronal transcriptional signatures^42^. Our neighborhood maps now provides a joint spatial-molecular basis for the hypothesis that these neuronal, immune and glial responses are coupled through local genome-maintenance niches.

Vascular imaging provides a complementary anatomical foundation for our model. Bennett et al. (2024) documented altered vessel geometry and regional reductions in vascular and pericyte density, with prominent involvement of deep cortical layers, the hippocampal network and basal forebrain^13^. Our distance-resolved analysis links altered vascular cell DDR states to proximal vascular neighbors, intermediate immune neighborhoods and more distal oligodendroglial associations. In cortex and cerebral nuclei, vascular and oligodendroglial enrichment favored DDR-high vascular anchors, whereas immune enrichment generally favored DDR-low anchors. Endothelial ERCC1 deficiency was shown to disrupts blood–brain barrier function and altered endothelial sprouting^41^, supplying a mechanistic link between repair capacity and vascular remodeling. Together, these observations support the hypothesis that vascular genome maintenance can couple vessel integrity to immune access and oligodendroglial support, connecting regional vascular vulnerability to the surrounding tissue environment.

Two complementary spatial analyses made this connection measurable. Analysis of metacells linked neuronal DDR transcriptional states to local cell composition, and annuli then further resolved the identity and distance of neighbors around individual DDR-defined anchors. Their agreement connects population-level tissue composition with the organization of individual cellular neighborhoods. The radial analysis further separates proximal vascular associations from more distal oligodendroglial associations, defining distinct spatial readouts for vascular maintenance and parenchymal support. This approach gives the proposed connection between repair programs and tissue remodeling a measurable anatomical structure.

As a practical extension of the annular analysis, we projected the age-specific radial association scores onto a common anatomical framework to generate brain-wide maps of DDR-state-conditioned cellular neighborhoods (Fig. 8 and Extended Data Fig. 5). This representation is distinct from maps of DDR-module expression or coexpression (Fig. 2) and from cell-type abundance (Fig. 4): it shows whether a selected immune, oligodendroglial or vascular neighbor population is preferentially associated with DDR-low or DDR-high neuronal or vascular cell anchors in each region and age group. The cortical vascular–immune examples for BER and NER, cortical neuron–oligodendroglial relationships for DSB, and thalamic vascular–immune relationships for TLS illustrate how the direction and strength of such preferences differ with aging. These maps thus provide a novel visualization tool of the aging brain and a regional reference for prioritizing anchor–neighbor combinations for targeted spatial validation and for selecting microenvironmental readouts following genome-maintenance perturbations. In pathological contexts, such comparative maps drawn from aging-associated diseases and healthy controls can help highlight changes beyond gene expression and spatial cell type abundance and can be customized for any relevant molecular correlates.

**Figure 8.**
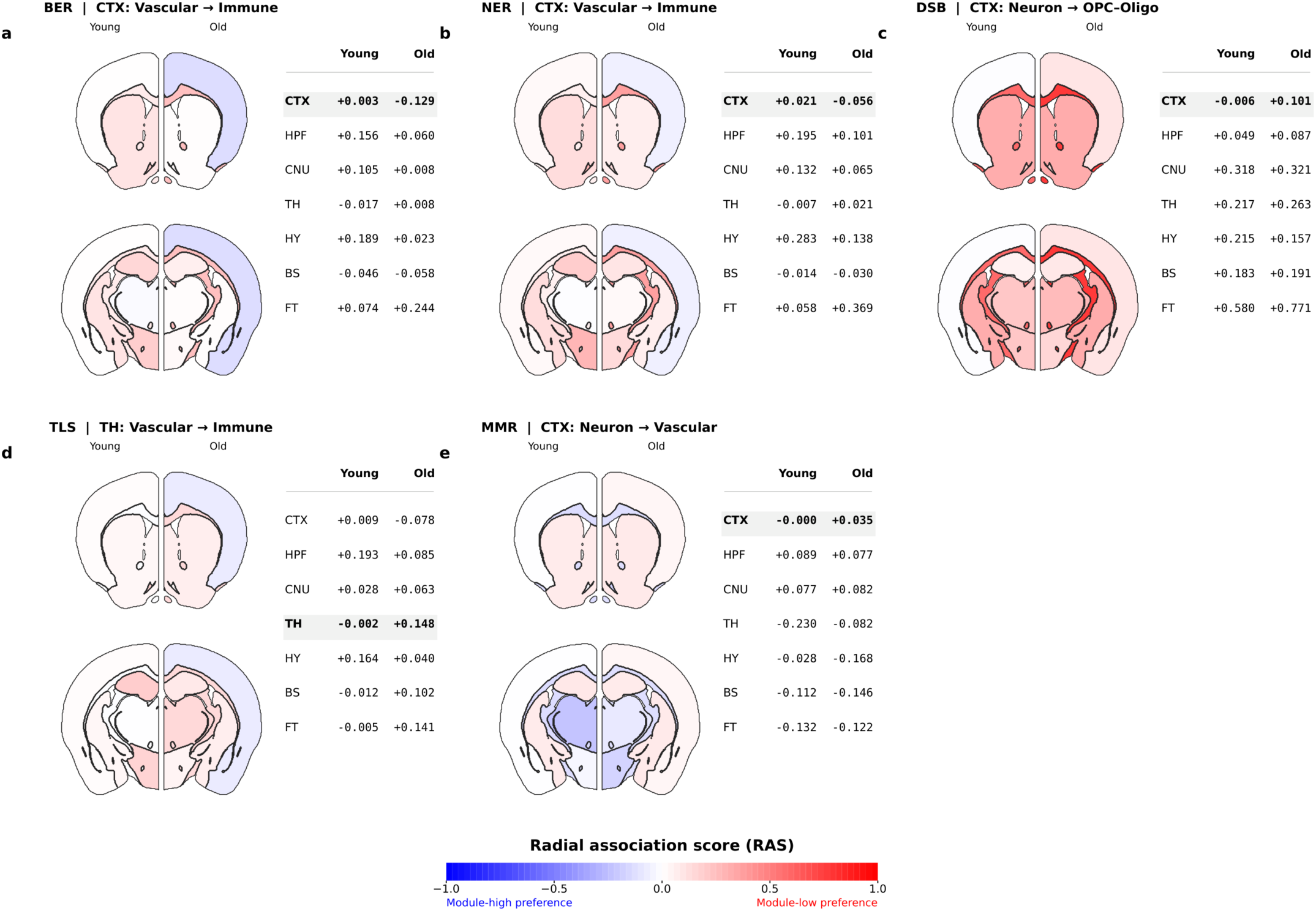
Anatomical maps of age-associated DDR-state-conditioned cellular microenvironments. A–E, Representative brain-wide maps of age-specific radial association scores (RAS) for BER vascular anchors and immune neighbors in cerebral cortex (A), NER vascular anchors and immune neighbors in cortex (B), DSB neuronal anchors and OPC–oligodendroglial neighbors in cortex (C), TLS vascular anchors and immune neighbors in thalamus (D), and MMR neuronal anchors and vascular neighbors in cortex (E). For each relationship, the left half of each coronal atlas representation displays Young and the right half Old. Both anterior and posterior reference sections show the same broad-region RAS assignments; the corresponding Young and Old numerical values for CTX, HPF, CNU, TH, HY, BS and FT are listed alongside, focal exemplar region emphasized. RAS is the annulus-width-weighted average of signed Cohen’s d across 0–150 μm, comparing the proportion of the indicated neighboring population around DDR-low versus DDR-high anchors within an age group. Blue (negative) denotes preferential representation around DDR-high anchors; red (positive) denotes preference around DDR-low anchors; color intensity represents the standardized difference rather than cell number or absolute abundance. Regional scores are applied to atlas territories for visualization and do not indicate within-region fine-scale variation or independent statistical significance. The complete six-module atlas is shown in Extended Data Fig. 5.

The oligodendroglial relationships suggest distinct routes through which genome maintenance engages tissue support. It was shown that cerebral endothelial trophic factors can promote OPC survival and proliferation, while oxidative stress disrupts this support^43^. The enrichment of oligodendroglial neighbors around DDR-high vascular anchors supports the hypothesis that vascular repair capacity sustains the oligovascular niche. Oligodendroglial enrichment around DDR-low neurons points to a complementary response to neuronal maintenance demands, consistent with the metabolic support that oligodendroglia provide to axons^4^. We propose that these configurations represent vascular support of the oligodendroglial lineage and oligodendroglial support of stressed neuronal environments, placing repair capacity within the reciprocal maintenance of the neurovascular unit.

Experimental work also connects genome stress to local immune responses. DSB-bearing neurons can engage antiviral-like signaling, and neuronal NFκB knockdown reduces nearby microglial activation^33^. This mechanism supports the hypothesis that neuronal genome stress recruits inflammatory responses within the immune-rich neighborhoods identified here. APOE4-associated dysfunctional human microglia were identified within niches characterized by gliosis and senescence^44^. Their observations support a disease-relevant rationale of our model in which genetic susceptibility amplifies inflammatory stress within genome-maintenance niches. Together with experimental evidence linking endothelial repair to barrier function and microglial DNA sensing to neurological aging^41,45^, these findings connect neuronal and vascular maintenance to shared immune responses through distinct local routes.

Our model predicts that altering neuronal or vascular repair capacity will in turn remodel distinct immune and oligodendroglial neighborhoods. Neuron-restricted repair perturbations should change local immune and oligodendroglial proximity, whereas vascular repair perturbations should change proximal vascular organization, immune access and the more distal oligodendroglial environment. Reciprocal perturbation of immune or oligodendroglial populations will test their influence on anchor-cell repair programs. Combining these interventions with direct measurements of DNA lesions, repair activity and vascular function will identify the mechanisms linking molecular maintenance to neighborhood remodeling. Across regions, the model predicts that repair-program organization and local cellular context together will explain vulnerability more effectively than either feature alone. Beyond proposing a conceptual framework our approach offers an immediate practical application, mapping the aging brain in terms of DDR-module-associated microenvironmental remodeling. Applying these spatial analyses to normative and pathological aging, including Alzheimer’s disease, could identify shared and divergent neighborhood patterns. Appropriately powered, sex-balanced studies could likewise test for sex-associated differences in these relationships. Genome-maintenance niches thus provide a biological framework in which established neuronal, immune, glial and vascular features of aging converge, with repair capacity as a proposed regulator of their local relationships.

## Methods

### Animals, tissue preparation and MERFISH

All animal care and experimental procedures were approved by the Pennsylvania State University Institutional Animal Care and Use Committee (IACUC). Spatial transcriptomic data were generated from C57BL/6J mice comprising young (n = 3; 2–3 months; 2 males and 1 female) and old (n = 3; 22 months; 2 males and 1 female) groups. One young and one old brain were processed together during tissue preparation and cryosectioning. From each animal, one anterior hemicoronal section (approximately bregma +0.98 mm) and one posterior hemicoronal section (approximately bregma −1.06 mm) were analyzed, yielding 12 sections across six paired MERFISH runs.

Brains were rapidly dissected after cervical dislocation and decapitation, embedded in Optimal Cutting Temperature compound with paired young and old hemibrains anatomically aligned, rapidly frozen in dry-ice-cooled 2-methylbutane, and stored at −80°C. Coronal sections were collected at 10-µm thickness. Anatomical levels were verified during sectioning using DAPI-stained adjacent sections, and RNA integrity was assessed from neighboring sections before MERFISH processing; only samples with RNA integrity number (RIN) ≥7 were advanced.

MERFISH^18^ was performed on a Vizgen MERSCOPE M1 instrument using V1 chemistry and a custom 500-gene panel containing cell-identity markers and genes relevant to the biological programs examined here. MERSCOPE-mounted sections were fixed in 4% paraformaldehyde for 15 min, permeabilized in 70% ethanol overnight at 4°C, hybridized with the gene-panel probes for 36–48 h at 37°C, washed, hydrogel embedded, cleared with Proteinase K, stained with DAPI and poly(T), and imaged according to established MERSCOPE procedures and vendor guidance. The detailed tissue-processing and MERFISH workflow is provided in the Supplementary Methods such as fixation, hybridization, 47°C post-hybridization washes, hydrogel embedding, overnight clearing and DAPI/poly(T) labeling used for this study.

### Segmentation, cell annotation and anatomical registration

Watershed segmentations were refined using a custom Cellpose2 model^46^ trained with manually annotated boundaries; its estimated F1 score was approximately 0.90. We assigned cell annotations using MetaMarkers^19^ to an adult mouse-brain reference taxonomy^20^, using the top 50 AUROC-ranked markers per cell type at the class level. DAPI images and spatial metadata were aligned to Allen CCFv3^21^ using QuickNII^47^ and refined by nonlinear alignment in VisuAlign^48^. Registered atlas-label images were used to assign cells and transcripts to CCF-defined regions. Cells outside labeled regions and cells with 0 detected transcripts across all genes were excluded; cells with an “unassigned” MetaMarkers annotation were excluded from analyses requiring cell-type identity. We normalized gene counts within each cell to 1 across all genes to control for differences in transcript detection between cells. Sections were retained as section-level observations and linked to their animal of origin. Registration accuracy was assessed against the corpus callosum, olfactory tubercle and striatal landmarks and independently evaluated by two investigators. A region absent from a section was treated as missing. Distance-resolved neighborhoods were calculated from measured cell coordinates within each section; CCF labels stratified the anatomical analyses but did not generate cell-cell adjacency. Normalized values were interpreted as relative expression within the targeted panel, not absolute transcript abundance, and the two sections from one mouse did not increase animal-level n.

### DDR modules and regional expression-coexpression analysis

Genes were grouped into BER, NER, MMR, DSB, TLS and CoreRep modules according to established functional roles; composite DDR scores integrated the pathway modules used in each analysis by module-level mean. To find regional differential expression, we measured the average cell expression for each gene within each animal, region and cell type to produce per-gene animal-level pseudobulk estimates. We performed a permutation test on the young and old distributions of these pseudobulk estimates, using a two-sided empirical null from 10,000 random sign assignments with a +1 continuity correction. We performed this test separately by cell type and combined the resulting P values using Fisher’s method to find an overall regional summary. We applied Benjamini-Hochberg FDR correction across these resulting overall region P values. To find regional differential coexpression, we measured the average cell expression for each gene within each animal, region and cell type to produce per-gene animal-level pseudobulk estimates. We then computed Pearson correlations across these pseudobulk profiles within each age for every pair of genes within each gene module. Fisher z-transformed old and young correlations were compared across matched gene pairs by paired t-test. To find overall region-level coexpression differences, we additionally performed this t-test on pseudobulked expression profiles for all cells in each region, followed by FDR correction across these whole-region P values. Across all regions, we only included comparisons meeting the prespecified minimum of at least three animals per age; regions failing this criterion were excluded.

Inter-module analysis used 123 matched fine regions with complete values for all six modules as the paired inferential units. For each age, overall within-module coexpression means were correlated across regions. Age-associated changes in correlation were tested by 10,000 within-region young-old label swaps, and 95% confidence intervals were obtained from 10,000 paired regional bootstraps. Benjamini-Hochberg FDR correction was applied across the 15 module pairs; no age-difference edge survived this correction. Spearman correlations and residualization within major anatomical compartments were used as sensitivity analyses. The matched region was the permutation and bootstrap unit; neither cells nor gene pairs supplied additional replication. Pooled regional coexpression was interpreted as a tissue-state measure that can include cellular-composition effects, not as proof of within-cell-type regulatory coordination.

### Cellular composition

For each sample and region, cell-type composition was calculated as the proportion of all detected cells, including zero proportions when a sampled region lacked a given cell type. Old and young proportions were compared by two-sided Welch’s t-test; P values were FDR corrected across cell types within each region. To find these changes in cell-type composition in broader regions, we recomputed cell proportions in those larger regions and performed the same t-test. A region absent from a section was treated as missing, not as zero; proportions quantify relative representation among mapped cells rather than absolute cell density.

### Spatial metacells

Spatial metacells were generated independently within each section by k-means clustering of cellular coordinates, targeting approximately 100 cells per metacell. Neuronal expression was pseudobulk-averaged across all neuronal cells in each metacell. To score metacells by their expression for any gene set, we ranked by percentile all expressed genes in that metacell, excluding unexpressed genes. We then found the average rank of genes within that gene set amongst all expressed genes. This average rank is the gene-set score for a given metacell. Cell-type proportions were calculated independently among all cells in each metacell. Composite-DDR-ranked metacells were divided into 50 equal-rank groups. For individual curated or GO gene sets, target-cell proportions in the highest and lowest groups were compared by Welch’s t-test with Benjamini-Hochberg correction. Metacells remained nested within sections and animals, leaving biological replication unchanged. Their tests quantify within-cohort associations between local neuronal gene-set state and local composition. Rank-based scores capture relative deployment within the targeted panel while reducing dependence on total transcript detection; they do not measure absolute pathway activity.

### Annular neighborhood analysis and age-polarization score

Cell-level DDR module scores used the same rank-among-expressed procedure: expressed genes were percentile-ranked within each cell, unexpressed genes were excluded from the ranking, and the module score was calculated from the mean percentile rank of detected genes belonging to that module. For each module, anchor cells at or below the 6th percentile were classified as DDR-low and anchors at or above the 94th percentile as DDR-high. Surrounding cells were partitioned into ten non-overlapping annuli spanning 0–150 μm: 0–5, 5–10, 10–20, 20–30, 30–40, 40–50, 50–75, 75–100, 100–125 and 125–150 μm. LIANA^49^ defined spatial connectivity, and neighbor composition was calculated as the proportion of all cells in each annulus. Within age and region, DDR-low and DDR-high neighborhoods were compared by two-sided Mann-Whitney U test, with Cohen’s d oriented as DDR-low minus DDR-high. P values were Bonferroni-corrected within each anatomical comparison. These rank-based scores represent relative deployment of the measured DDR genes within the targeted panel; DNA-lesion burden and repair capacity require direct assays. Annuli were constructed from measured within-section coordinates, with CCF labels used only to stratify analyses by anatomical region. Anchor cells remained nested within sections and animals. The Mann-Whitney tests quantify anchor-level spatial association within the observed tissue, with animal-level sample size fixed at six. Spatial proximity and anchor-to-neighbor notation carry no implication of recruitment, signaling or causal direction.

For each age, all finite annulus-specific Cohen’s d values were multiplied by the corresponding annulus width, summed and divided by the total sampled radial width to yield the age-specific radial association score. No effect-size or significance filtering was applied before integration. The age-polarization score (APS) was defined as the old radial association score minus the young radial association score. Positive APS denotes a shift toward DDR-low-associated neighborhoods and negative APS a shift toward DDR-high-associated neighborhoods. APS is a descriptive difference between independently estimated age-specific profiles and was not treated as a formal age-by-DDR interaction test. Plot outlines denote significance in at least one contributing annulus; stars encode the maximum absolute annulus-level effect size and are not significance indicators. Arrows between young and old scores visualize direction of chronological aging between descriptive summaries and do not imply the direction of signaling.

### Statistical units and scope of inference

Analytical units were matched to the question and are stated above. Large cell numbers improve estimation of animal-level pseudobulk values or local spatial distributions but do not increase the six-animal biological cohort. Animal-level regional estimation, matched-region inter-module inference and nested metacell or anchor-level association were therefore interpreted separately. Concordance between metacell and annular representations was treated as within-cohort convergence, not as orthogonal experimental validation. All DDR-high and DDR-low relationships reported here are cross-sectional associations and do not establish causal or signaling direction.

## Acknowledgements

We thank the Strategic Instrumentation and Pilot Projects mechanism at Penn State College of Medicine for the acquisition of MERSCOPE equipment. We thank members of the Paul and Kim laboratory for helpful discussions and feedback throughout the project. Additionally, we thank the high-performance computing (HPC) center and data storage at The Penn State College of Medicine.

## Author contributions

**Conceptualization:** Anirban Paul

**Data curation:** Steffy B. Manjila, Mofida Abdelmageed

**Formal analysis:** Arvin Khoshboresh, Vijay Laxmi Roy

**Funding acquisition:** Anirban Paul

**Investigation:** Arvin Khoshboresh, Vijay Laxmi Roy, Anirban Paul, Jesse Gillis

**Methodology:** Arvin Khoshboresh, Vijay Laxmi Roy, Steffy B. Manjila, Mofida Abdelmageed, Deniz Parmaksiz, Yongsoo Kim, Anirban Paul, Jesse Gillis

**Project administration:** Anirban Paul

**Resources:** Anirban Paul, Jesse Gillis, Yongsoo Kim

**Software:** Arvin Khoshboresh, Vijay Laxmi Roy, Deniz Parmaksiz

**Supervision:** Anirban Paul, Jesse Gillis

**Visualization:** Arvin Khoshboresh, Vijay Laxmi Roy, Anirban Paul

**Writing – original draft:** Anirban Paul

**Writing – review & editing:** Anirban Paul, Arvin Khoshboresh, Jesse Gillis, Vijay Laxmi Roy, Yongsoo Kim, Steffy B. Manjila, Deniz Parmaksiz

## Availability of data and materials

The code, processed data, and analysis scripts supporting the findings of this study will be made publicly available upon publication. The primary MERFISH dataset will be deposited in a public repository, and source code required to reproduce the analyses, including the spatial transcriptomics analysis pipeline and example workflows, will be made available through GitHub with archived releases assigned DOIs. Repository accession numbers and URLs will be added prior to publication.

## Ethics approval and consent to participate

Not applicable.

## Consent for publication

Not applicable.

---

**Extended Data Figure 1.**
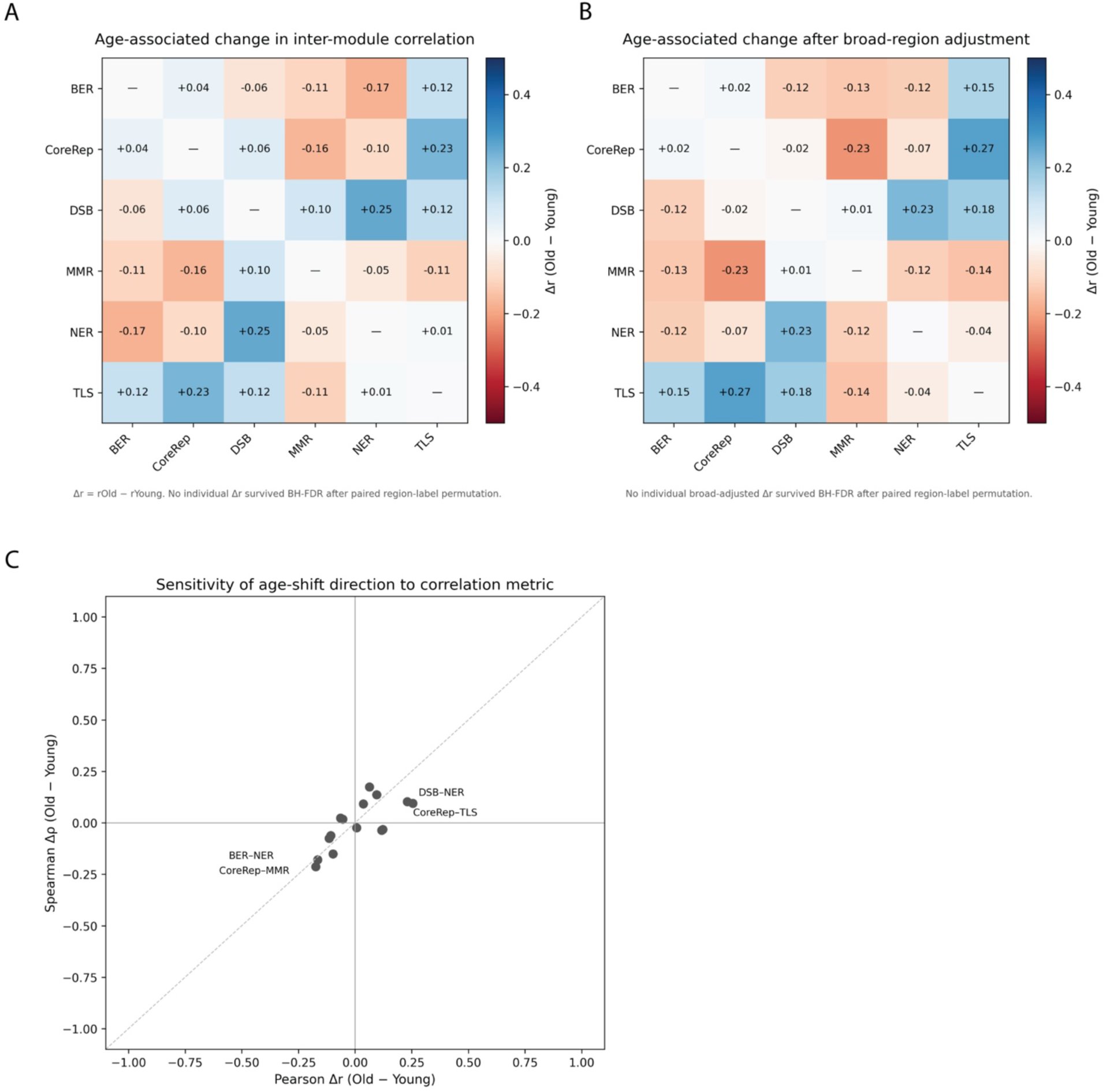
Age-associated changes in inter-module coexpression among DNA damage response programs across matched brain regions. A, Heat map of the old-minus-young change in pairwise inter-module Pearson correlation (Δr) across matched fine brain regions. Positive values indicate stronger and negative values weaker inter-module covariance in old brain. **B,** The same analysis after removal of broad-region effects. **C,** Sensitivity analysis comparing Pearson Δr with Spearman Δρ for each module pair; labeled points identify representative relationships. The direction of age-related change was largely preserved across correlation metrics. No individual unadjusted or broad-region-adjusted age difference survived BH-FDR correction using paired region-label permutation testing.

**Extended Data Figure 2.**
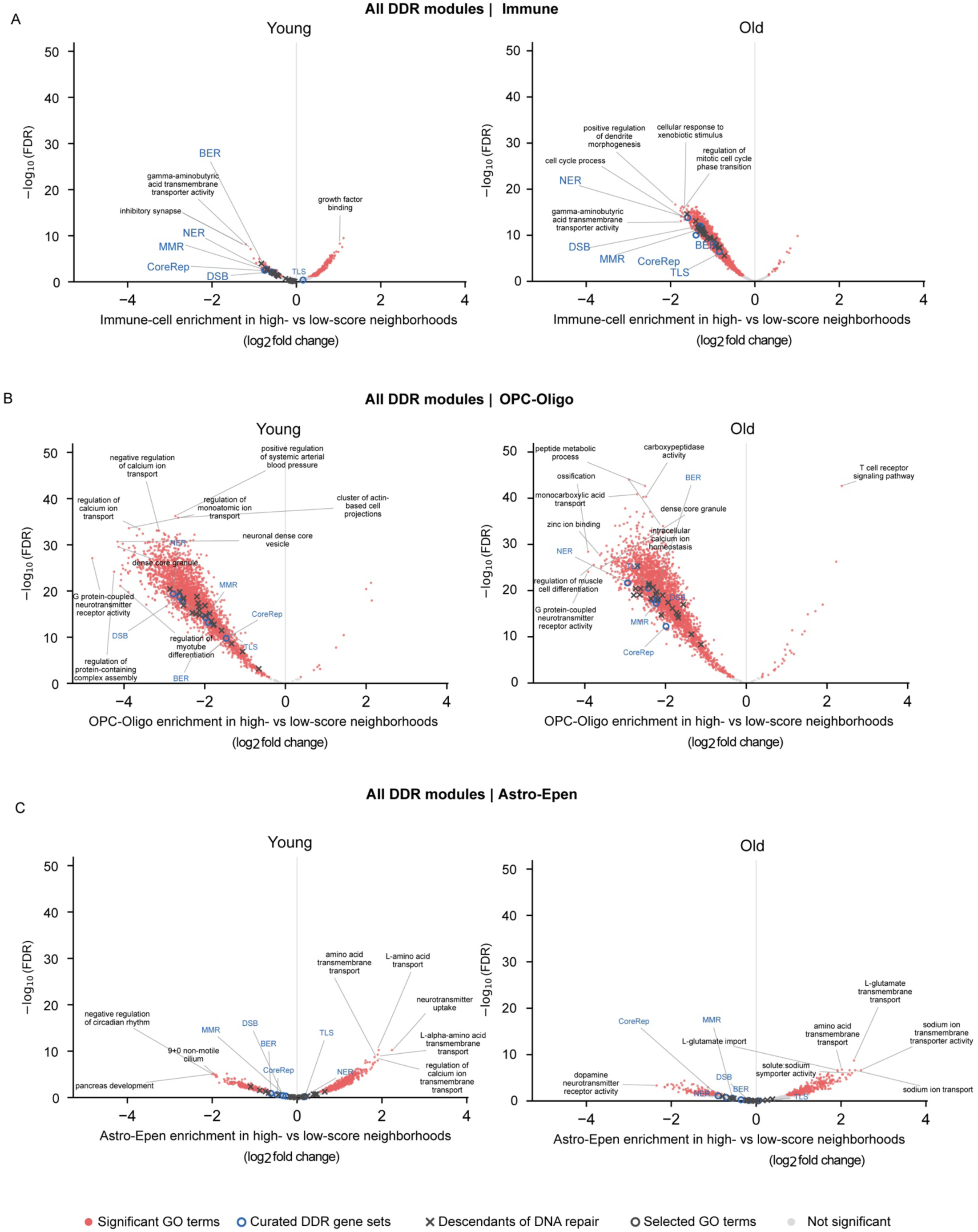
Gene-set-level associations between neuronal pathway state and local non-neuronal composition. A-C, Gene-set-level association between neuronal pathway scores and local abundance of immune (A), OPC-Oligo (B) and Astro-Epen (C) cells in young (left) and old (right) brains. Metacells were independently ranked for each of approximately 4,000 GO terms, curated DDR gene sets and descendants of the GO DNA-repair term, and target-cell proportions were compared between the highest- and lowest-scoring bins. The x axis shows log2 fold change in target-cell abundance in high-versus low-scoring neuronal metacells; negative values therefore indicate enrichment of the indicated cell population in low-scoring neuronal neighborhoods, whereas positive values indicate enrichment in high-scoring neighborhoods. The y axis shows -log10(FDR). Significant GO terms are shown in red, curated DDR gene sets as blue open circles, descendants of DNA repair as black crosses, selected GO terms as black open circles and non-significant terms in grey. Two-sided Welch’s t-tests were used to compare the highest- and lowest-scoring bins, with Benjamini–Hochberg FDR correction across the tested gene sets. Effect sizes and the direction of the DDR-ranked gradients were interpreted together with multiplicity-corrected P values. Abbreviations: Astro-Epen, astrocyte–ependymal; BER, base excision repair; CoreRep, core replication; DDR, DNA damage response; DSB, double-strand break repair; Epen, ependymal; FDR, false discovery rate; GO, Gene Ontology; MMR, mismatch repair; NER, nucleotide excision repair; OPC, oligodendrocyte precursor cell; OPC-Oligo, oligodendrocyte precursor–oligodendrocyte; TLS, translesion synthesis.

**Extended Data Figure 3.**
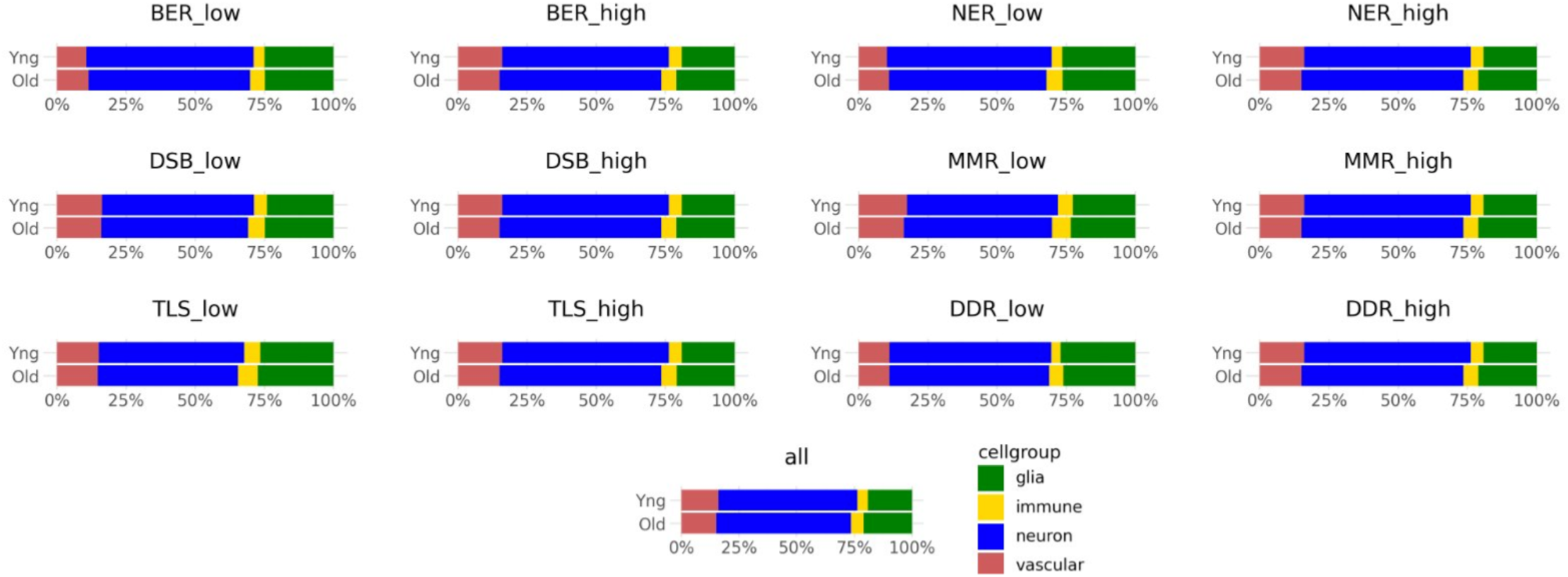
Broad cell-class composition of DDR-low and DDR-high cell subsets in young and old brains. Stacked bars show the relative representation of four broad cell groups (glia, immune, neuron and vascular) among all mapped cells (all) and among cells in the low and high extremes of BER, NER, DSB, MMR, TLS and composite DDR scores, shown separately for young and old samples. Each bar is normalized to 100% within the indicated age and DDR-defined subset, and colors denote broad cell group. Across individual modules and composite DDR, the DDR-low and DDR-high pools retained similar broad cell-class composition within each age and broadly recapitulated the overall cellular mixture. These bars are descriptive pooled-cell summaries and do not define biological n. This compositional control argues against a gross imbalance in neuronal, glial, immune or vascular representation among DDR-defined populations as a trivial explanation for the anchor-centered neighbor-composition differences in Figs. 6 and 7. The distance-resolved analyses therefore condition on neuronal or vascular anchor identity and quantify spatial organization of neighboring cells rather than simply reflecting a different broad cell-class mixture in the DDR-low and DDR-high pools. Abbreviations: BER, base excision repair; DDR, DNA damage response; DSB, double-strand break repair; MMR, mismatch repair; NER, nucleotide excision repair; TLS, translesion synthesis.

**Extended Data Figure 4.**
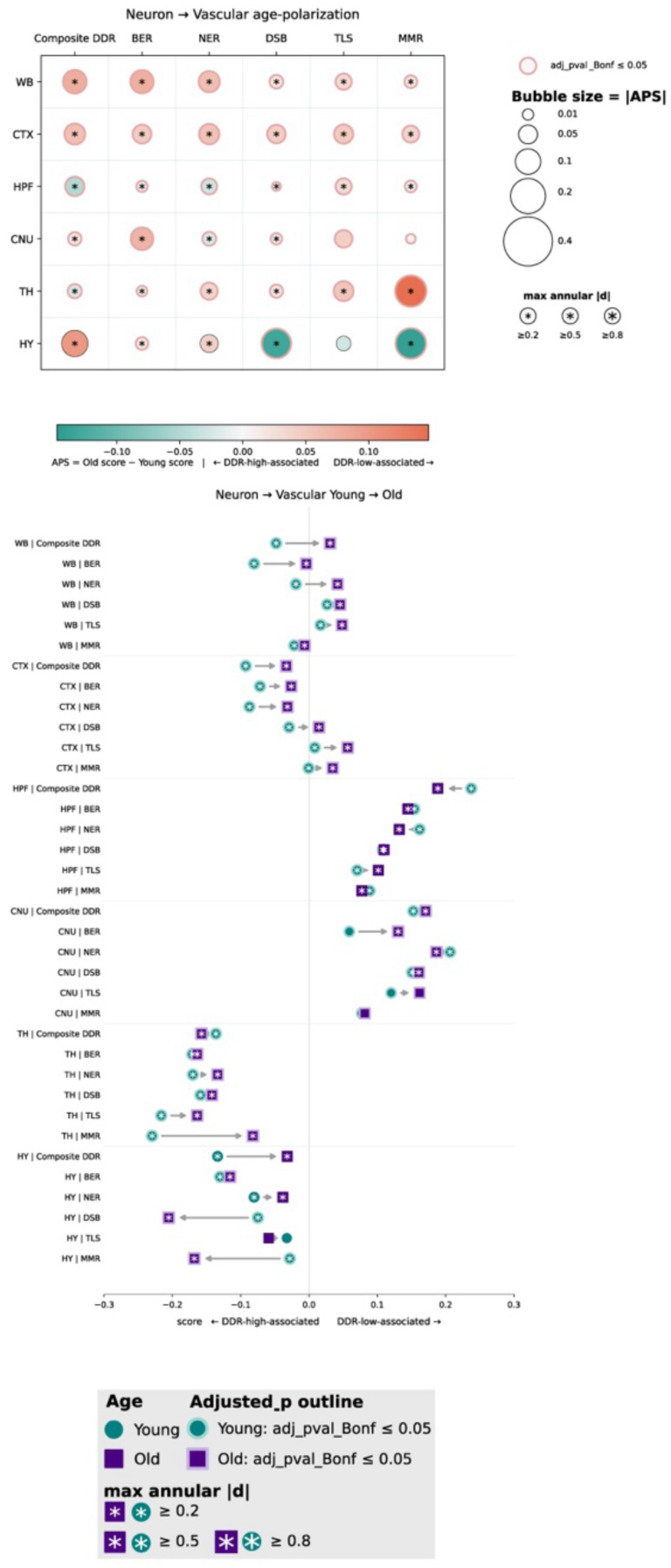
Age-dependent polarization of vascular neighborhoods surrounding DDR-defined neuronal states A, Brain-wide summary of age-dependent polarization of vascular-cell neighborhoods surrounding neuronal anchors. For each brain region and DDR module, annulus-level Cohen’s d values comparing vascular-neighbor abundance around DDR-low versus DDR-high neuronal anchors were integrated across the 0–150 μm radial neighborhood to yield an age-specific radial association score. The age-polarization score (APS) was calculated as the old radial association score minus the young radial association score. Positive APS values indicate an age-associated shift toward preferential vascular association with DDR-low neuronal anchors, whereas negative values indicate a shift toward DDR-high neuronal anchors. Bubble area is proportional to |APS| and bubble color denotes the direction of the age-associated shift. Outlines indicate Bonferroni-adjusted significance in the underlying annulus-level comparisons, and stars indicate the maximum absolute annulus-level Cohen’s d, with thresholds of |d| ≥ 0.2, ≥ 0.5 and ≥ 0.8. Columns show Composite DDR, BER, NER, DSB, TLS and MMR across WB, CTX, HPF, CNU, TH and HY. **B,** Directional young-to-old changes in neuron-to-vascular radial association. Arrows visualize direction of chronological aging between descriptive summaries and do not imply the direction of signaling. Teal circles and purple squares indicate young and old age-specific radial association scores, respectively, and arrows connect the scores for each brain region and DDR module. Negative scores indicate preferential vascular representation around DDR-high neuronal anchors, whereas positive scores indicate preferential representation around DDR-low neuronal anchors. Rightward arrows therefore indicate an age-associated shift toward DDR-low-associated vascular neighborhoods and leftward arrows indicate a shift toward DDR-high-associated neighborhoods; trajectories crossing zero denote reversal of the predominant DDR-state association with age. Marker outlines indicate Bonferroni-adjusted significance in at least one underlying annulus, and stars denote the maximum absolute annulus-level Cohen’s d using the thresholds shown. Age-specific radial association scores and APS were calculated from the unfiltered annulus-level effect-size profiles; the directional plots preserve the young and old spatial states, whereas APS summarizes their difference. The analysis comprised anterior and posterior coronal sections from three animals per age group (six sections per age group). Abbreviations: APS, age-polarization score; BER, base excision repair; CNU, cerebral nuclei; CTX, cerebral cortex; DDR, DNA damage response; DSB, double-strand break repair; HPF, hippocampal formation; HY, hypothalamus; MMR, mismatch repair; NER, nucleotide excision repair; TH, thalamus; TLS, translesion synthesis; WB, whole brain.

**Extended Data Figure 5.**
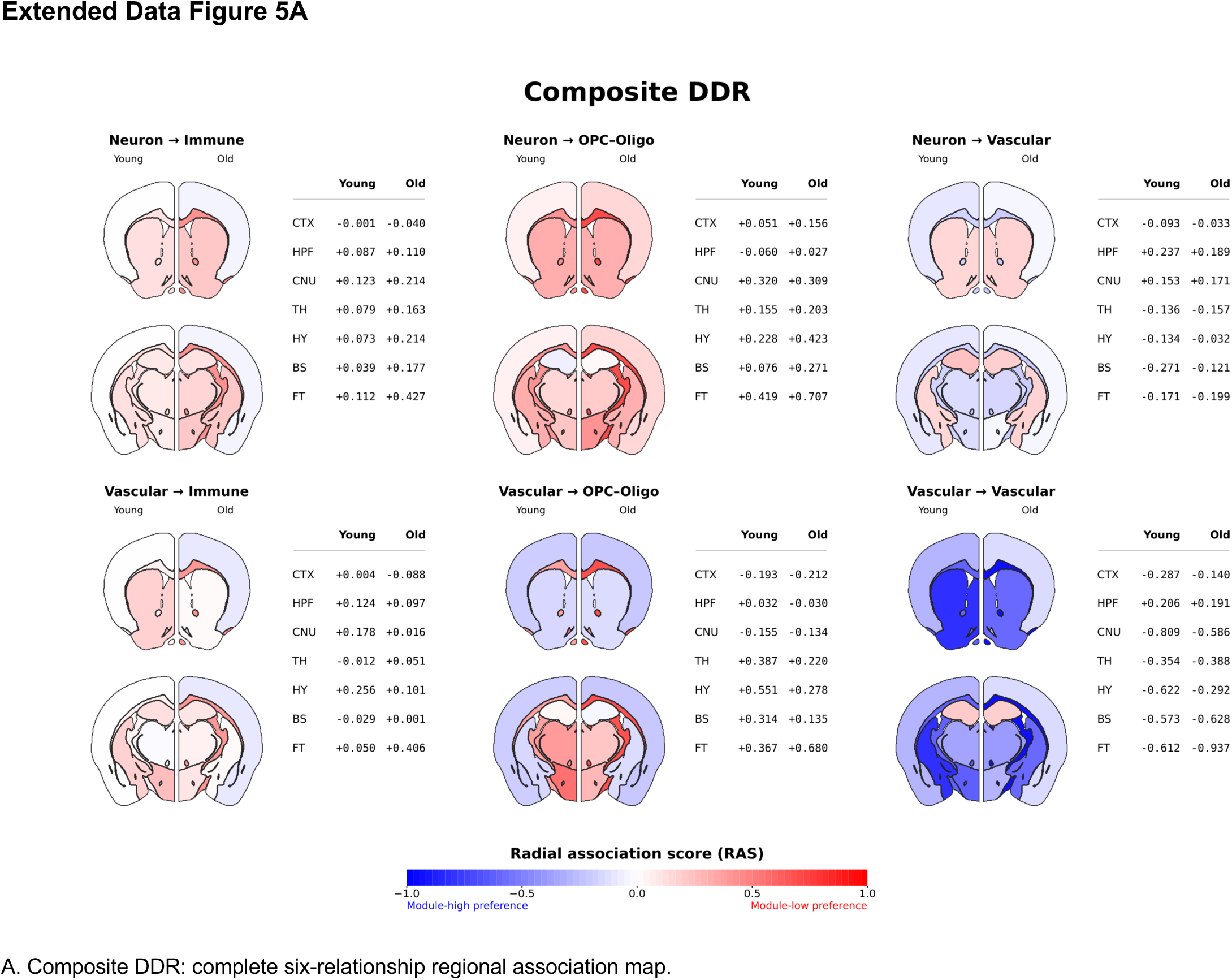
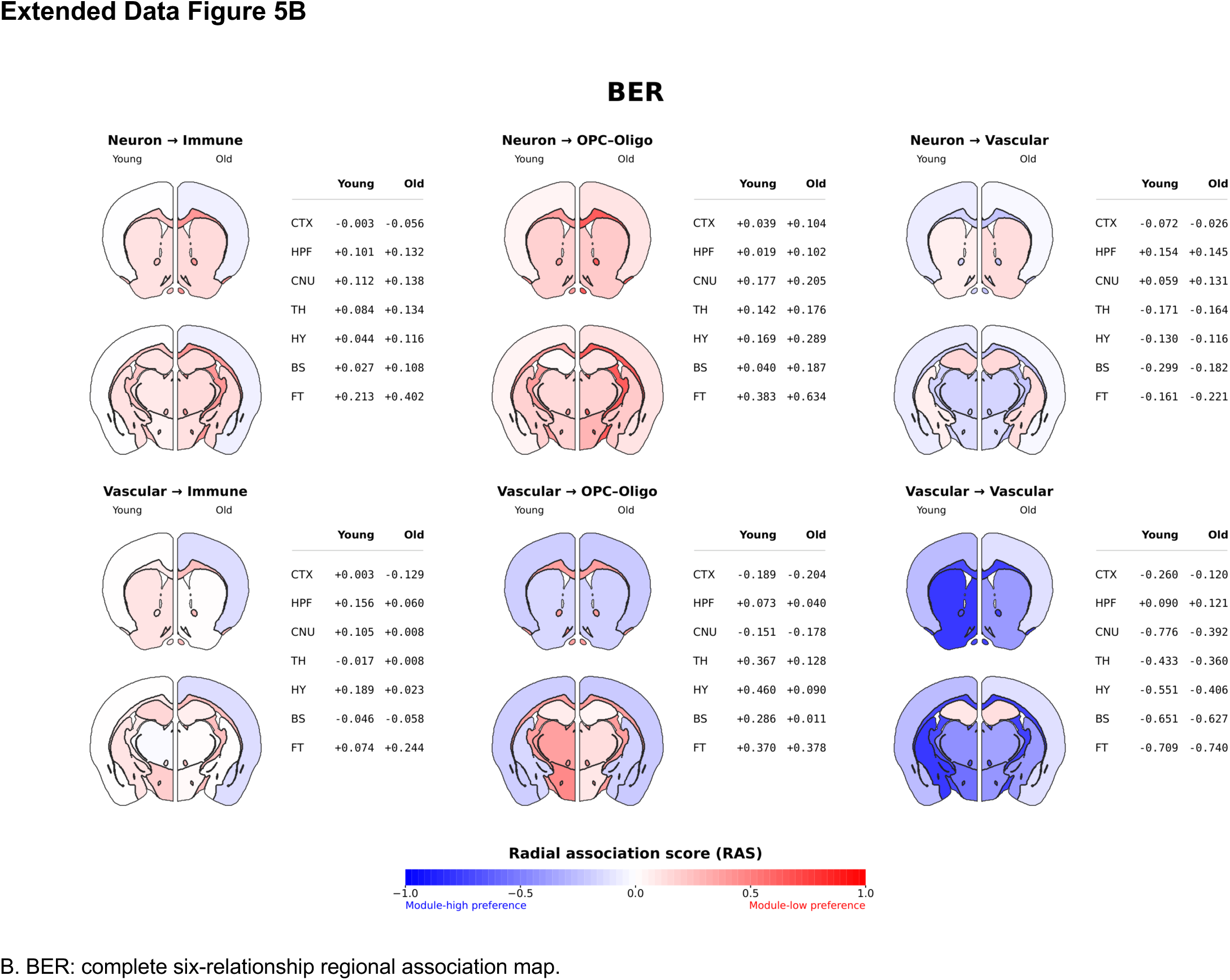
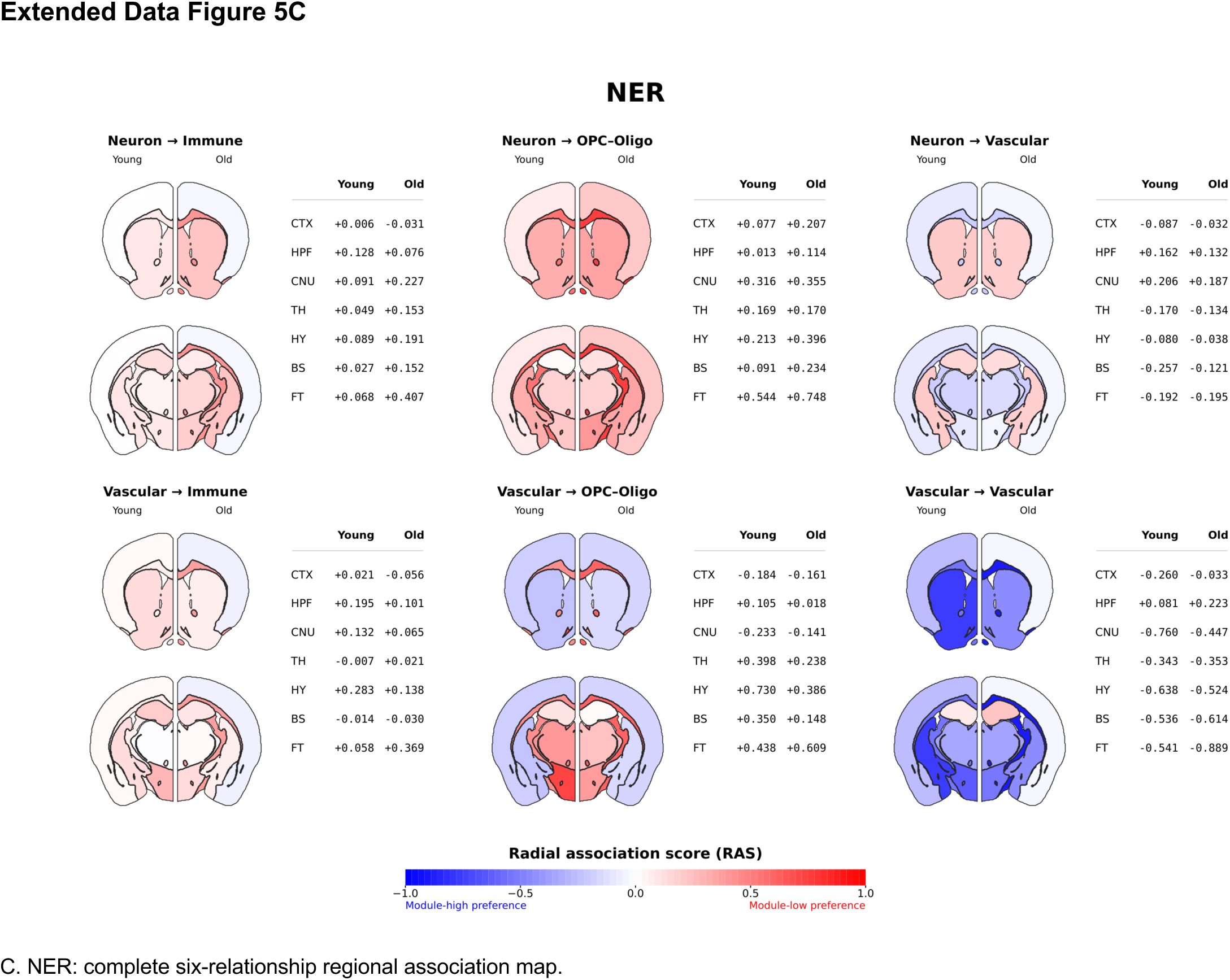
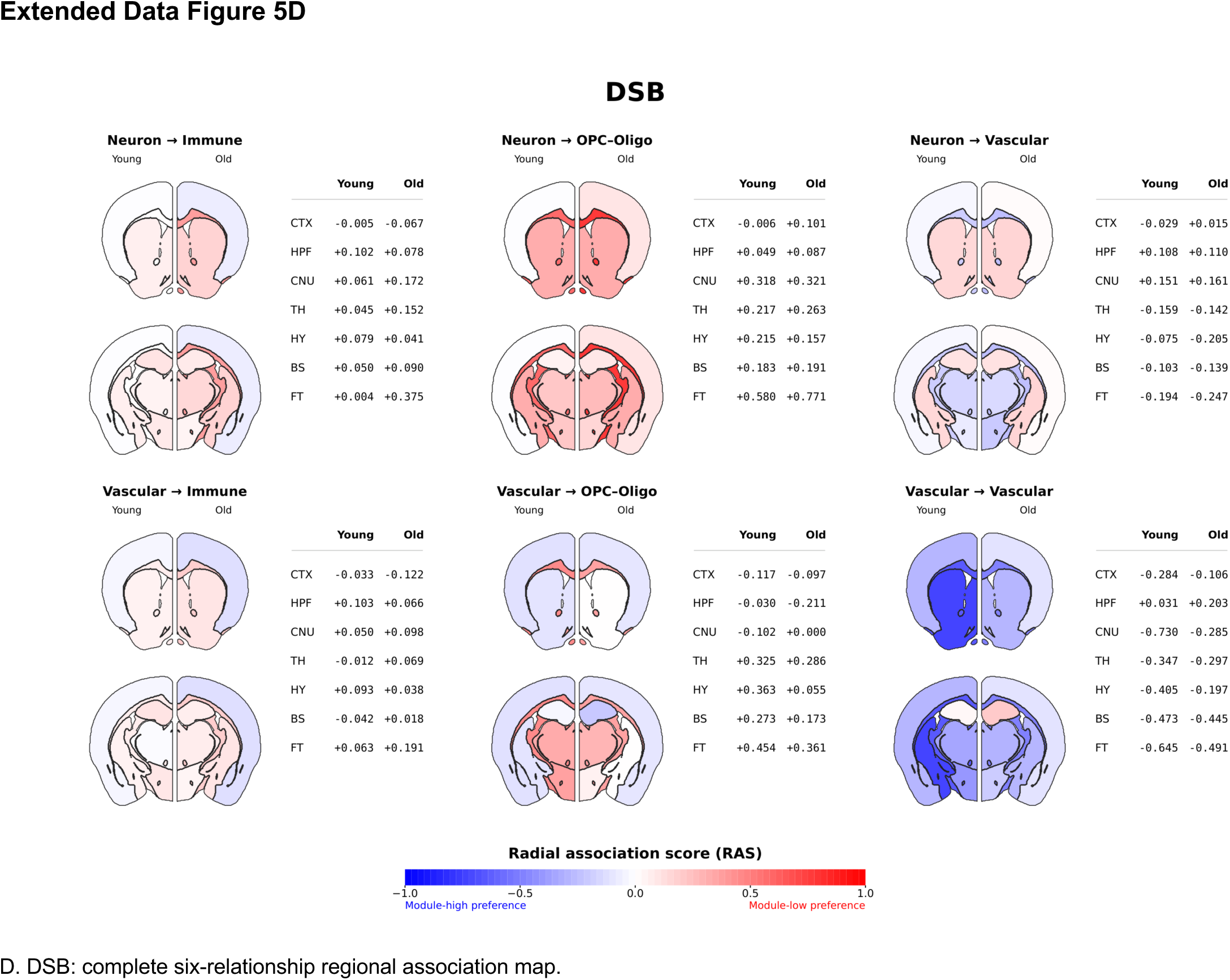
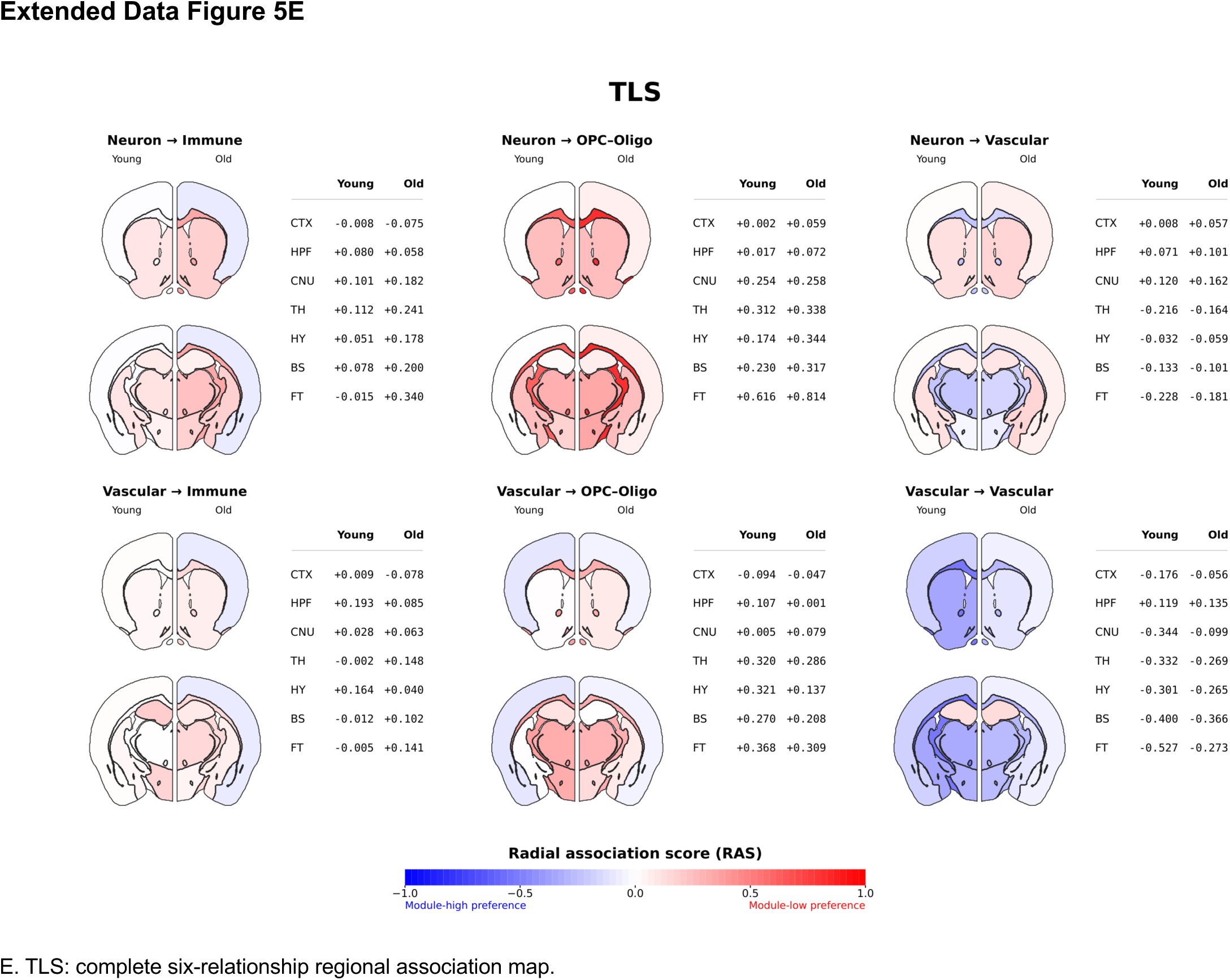
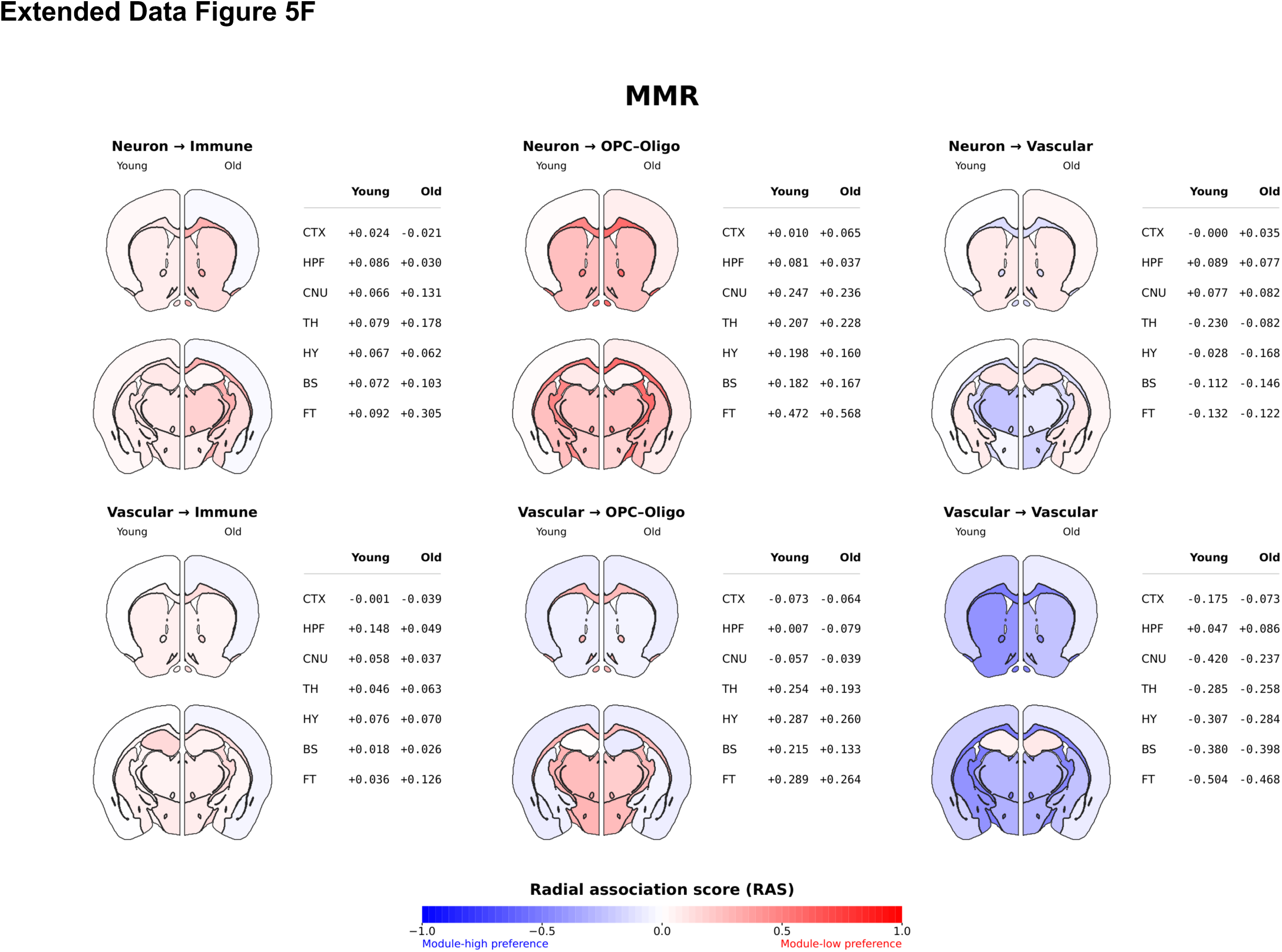
Complete Young–Old brain-wide atlas of DDR-state-conditioned cellular neighborhoods. A–F, Composite DDR (A), BER (B), NER (C), DSB (D), TLS (E) and MMR (F). Each module is shown for six anchor→neighbor pairs: neuronal anchors with immune, OPC–oligodendroglial or vascular neighbors (top row), and vascular anchors with the same three neighboring classes (bottom row). Left hemispheres represent Young and right hemispheres Old at two representative reference atlas levels. Numerical Young and Old radial association scores (RAS) are provided for CTX, HPF, CNU, TH, HY, BS and FT. RAS is the annulus-width-weighted mean of signed Cohen’s d across ten non-overlapping annuli spanning 0–150 μm, with Cohen’s d oriented DDR-low minus DDR-high. Blue (negative) indicates greater relative neighbor representation around DDR-high anchors and red (positive) around DDR-low anchors, using an identical −1 to +1 scale across all modules. Changes in hue or color intensity represent changes in the relative spatial association and do not, by themselves, establish changes in cell abundance, DDR expression or repair activity. Age-polarization scores (APS = Old RAS − Young RAS) describe change between the two age-specific scores; no independent significance test of APS is implied by the maps. Spatial assignments use broad anatomical regions rather than observed local spatial patterns at each point of the reference sections. These maps expand the representative examples in Fig. 8 and complement the distance-resolved analyses in Figs. 6 and 7. Abbreviations: APS, age-polarization score; BER, base excision repair; BS, brainstem; CNU, cerebral nuclei; CTX, cerebral cortex; DDR, DNA damage response; DSB, double-strand break repair; FT, fiber tracts; HPF, hippocampal formation; HY, hypothalamus; MMR, mismatch repair; NER, nucleotide excision repair; OPC–Oligo, oligodendrocyte precursor cell–oligodendrocyte; RAS, radial association score; TH, thalamus; TLS, translesion synthesis.

**Supplementary Figure 6.**
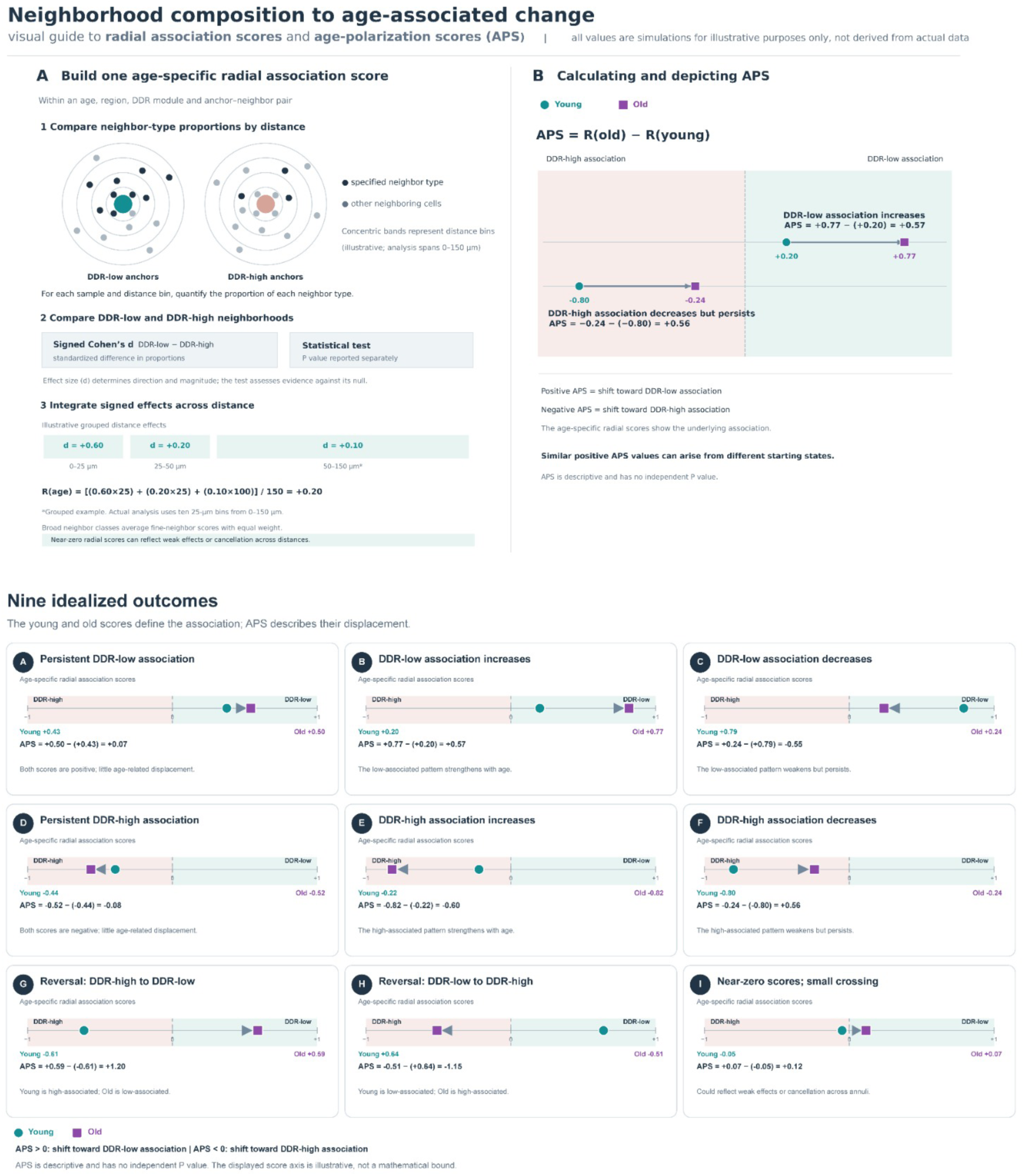
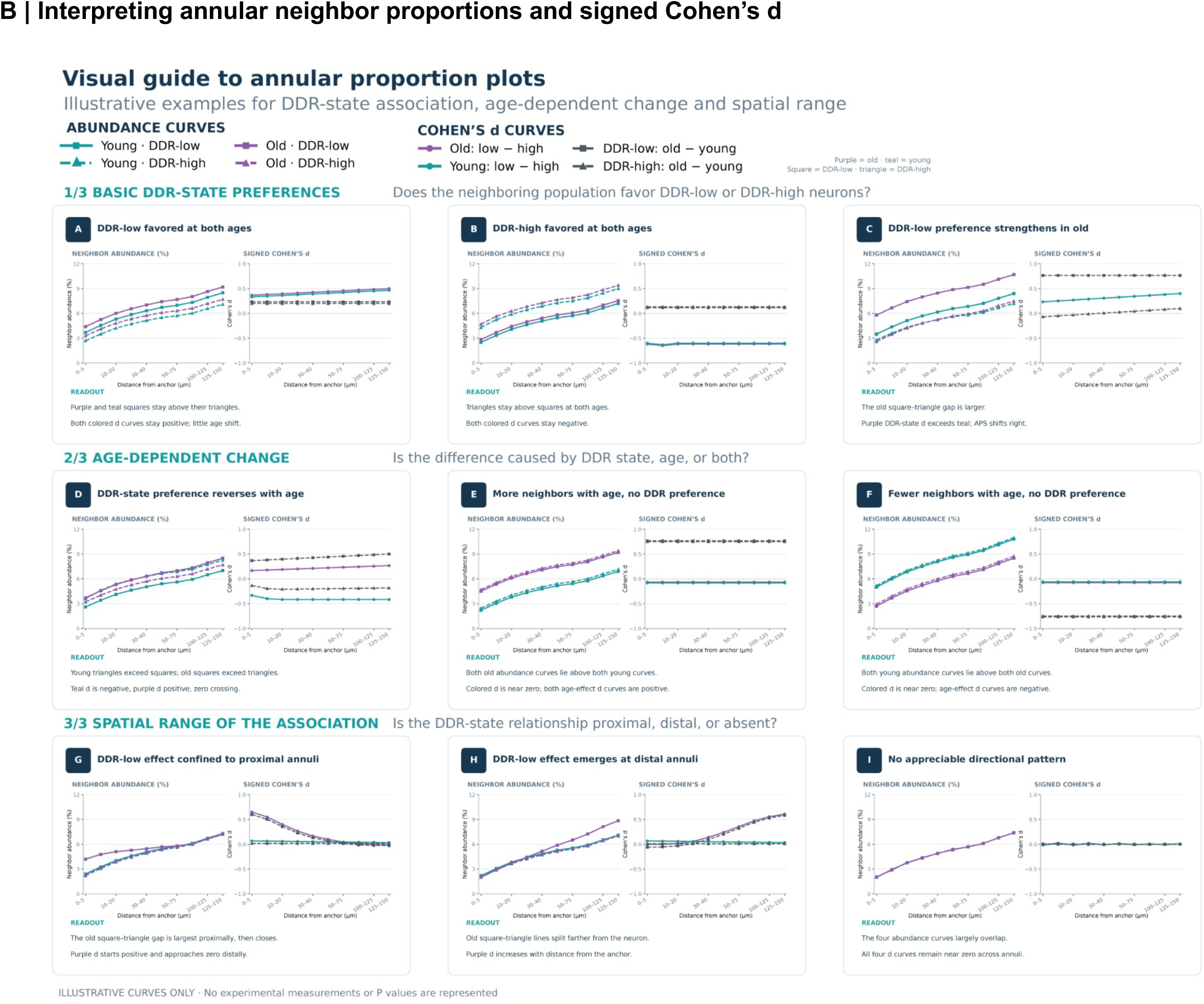
Visual guide to DDR-associated annular neighborhood analysis and age-dependent spatial associations. A, Schematic of how neighbor-type proportions are compared around DDR-low and DDR-high anchors across radial annuli, summarized as an annulus-width-weighted radial association score (RAS) and an age-polarization score (APS = Old RAS − Young RAS). Nine hypothetical score trajectories illustrate persistence, strengthening, attenuation, reversal and small near-zero crossings. B, Illustrative paired neighbor-abundance and Cohen’s d profiles distinguish DDR-low and DDR-high preference, age-associated abundance changes without a state preference, and proximal versus distal differences. Positive signed Cohen’s d favors DDR-low anchors, whereas negative values favor DDR-high anchors. *Both panels contain hypothetical examples solely for interpretation of* Figure 6 *and 7; they are not study measurements*.

## Supplementary Methods

### Animals and paired tissue collection

All animal care and experimental procedures were approved by the Pennsylvania State University Institutional Animal Care and Use Committee (IACUC). Experiments used C57BL/6J mice from two age groups: young mice (n = 3; 2–3 months old; 2 males and 1 female) and old mice (n = 3; 22 months old; 2 males and 1 female). One young and one old mouse were processed together during each tissue-preparation and cryosectioning session to minimize differences associated with sample handling. From each mouse, tissue was collected at two hemicoronal levels corresponding approximately to bregma +0.98 mm anteriorly and −1.06 mm posteriorly. The dataset comprised 12 sections from six biological replicates.

Mice were euthanized, immediately decapitated, and brains were rapidly dissected under RNase-free conditions. The first dissected brain was maintained on ice while the paired animal was processed, with extraction of each young–old pair completed within approximately 5–8 min. To maintain sample identity during co-processing, the young brain was divided in a sagittal plane slightly lateral to the midline, whereas the old brain was divided at the midline. Dissection instruments and blades were treated with RNaseZap (Invitrogen, AM9780) before and after use.

### Co-embedding, cryosectioning and RNA-quality assessment

Optimal Cutting Temperature compound (OCT; Tissue-Tek, 4853) was added to approximately one-fifth of a cryomold (Ted Pella, 27110). Paired young and old hemibrains were placed adjacent to one another in the same mold and aligned using the olfactory bulb, cerebral cortex and cerebellum as anatomical landmarks. The anterior–posterior orientation was recorded before the mold was completely filled with OCT. Blocks were rapidly frozen in 2-methylbutane pre-cooled with dry ice and stored at −80°C until sectioning.

Coronal sections were cut at 10-µm thickness in the anterior-to-posterior direction beginning at the olfactory bulb. During sectioning, mediolateral and anteroposterior tilt were adjusted as needed to obtain anatomically matched sections from the paired young and old hemibrains. As the target level was approached, approximately every tenth section was collected onto a conventional glass slide and stained with DAPI. Anatomical landmarks in these sections were used to confirm anatomical matching of the paired brains and proximity to the target bregma coordinates before tissue was collected onto MERSCOPE slides.

Following anatomical confirmation, 3–4 consecutive sections were collected onto MERSCOPE slides (Vizgen, 2040000), with the tissue positioned within the available imaging area. The present study analyzed one anterior and one posterior section from each animal. Approximately ten immediately adjacent sections were subsequently pooled and stored at −80°C for RNA-quality assessment. RNA was isolated using the RNeasy Mini Kit (Qiagen, 74104), and RNA integrity was assessed using an Agilent 2100 Bioanalyzer with an RNA Nano chip. Only tissue with RIN ≥7 was advanced for MERFISH analysis. Section identification and RNA-quality assessment were performed independently at the anterior and posterior sampling planes.

### MERFISH sample preparation and MERSCOPE imaging

MERFISH experiments were performed using the Vizgen MERSCOPE platform (M1 instrument, V1 chemistry) with a custom 500-gene panel containing cell-type markers and genes selected for the biological pathways analyzed in this study. Sample preparation followed the established fixed-frozen MERSCOPE workflow according to the manufacturer’s protocol for V1 chemistry.

Fresh-frozen 10-µm sections mounted on MERSCOPE slides were fixed in 4% paraformaldehyde in 1× PBS for 15 min, washed three times in 1× PBS and permeabilized in 70% ethanol at 4°C overnight. Sections were hybridized with the custom MERSCOPE Gene Panel Mix at 37°C for 36–48 h. Following probe hybridization, sections were washed twice in Formamide Wash Buffer for 30 min at 47°C. These fixation, permeabilization and hybridization conditions followed the Vizgen’s published protocols for V1 chemistry.

Hybridized sections were embedded in a hydrogel using Gel Embedding Premix (Vizgen, 20300004), ammonium persulfate (Sigma, 09913-100G) and TEMED (Sigma, T7024-25ML) from the MERSCOPE Sample Prep Kit. Hydrogel polymerization proceeded for approximately 1.5 h. Sections were then cleared overnight at 37°C in clearing solution containing Proteinase K (NEB, P8107S) and Clearing Premix (Vizgen, 20300003).

After clearing, sections were stained with DAPI and Poly T Reagent (Vizgen, 20300021) for 15 min at room temperature and washed for 10 min in Formamide Wash Buffer. Slides were assembled and imaged using the MERSCOPE system according to the manufacturer’s acquisition workflow.

The 12 analyzed sections were processed across six MERFISH imaging runs, with each run containing a matched young and old hemicoronal section. Anterior sections were processed in three runs and posterior sections in three runs, with the same biological pairs analyzed at both anatomical levels.

### Gene panel composition

The custom MERFISH gene panel was designed to support simultaneous identification of major brain cell populations and measurement of DNA damage repair (DDR)-related transcriptional programs. The panel incorporated a broad set of cell-type marker genes together with genes representing DNA repair and other biological processes. Cell-type markers were selected to distinguish major neuronal and non-neuronal populations and resolve neuronal subclasses and regional identities. Neuronal markers included genes distinguishing excitatory and inhibitory populations (for example, Slc17a6, Slc17a7, Slc17a8 and Slc32a1), major interneuron classes (Pvalb, Sst, Vip, Htr3a, Tac1, Chodl, Nos1 and Cck), and cortical projection-neuron and laminar identities (Cux2, Rorb, Tle4 and Tbr1). The panel also contained markers for oligodendrocyte-lineage cells and OPCs (Pdgfra, Cspg4, Cnp, Mobp, Mog and Cldn11), astrocytes (Aqp4, Aldh1l1, Fgfr3 and Mlc1), and immune/microglial populations (Csf1r, Cx3cr1, C1qa and Cd68). Panel included markers of cerebrovascular populations, including endothelial cells (Cldn5, Mfsd2a, Vwf and Pecam1), perivascular/mural populations (Pdgfrb, Kcnj8, Cspg4 and Anpep) and vascular smooth-muscle cells (Acta2, Tagln, Myh11, Cnn1 and Myl9). Additional markers distinguished selected cortical, hypothalamic neuronal populations. These markers supported broad cell-class assignment and subsequent spatial neighborhood analyses.

For genome-maintenance analyses, genes in the panel were grouped into six functional modules: base excision repair (BER), nucleotide excision repair (NER), mismatch repair (MMR), double-strand break repair (DSB), translesion synthesis (TLS) and CoreRep. Module assignments were based on established functional roles of the encoded proteins rather than on expression patterns observed in the present dataset. BER encompassed factors involved in recognition and repair of small base lesions and associated repair synthesis; NER included factors responsible for recognition and removal of bulky or helix-distorting DNA lesions; MMR comprised factors involved in mismatch recognition and processing; DSB included components of the principal pathways responsible for repairing DNA double-strand breaks; TLS comprised specialized DNA polymerases and associated factors that permit DNA synthesis across damaged templates; and CoreRep represented the core DNA-replication-associated program included in the targeted panel. Because several genome-maintenance proteins participate in more than one process, genes were assigned to the principal pathway used for module-level analyses without further subdivision into mechanistic submodules.

Genes were grouped into BER, NER, MMR, DSB, TLS and CoreRep modules according to established functional roles; Composite DDR scores were calculated as the mean of the pathway-module scores included in each analysis. Regional expression/coexpression, inter-module and metacell analyses used the six-module framework. The annular neighborhood analyses used BER, NER, MMR, DSB and TLS together with a Composite-DDR score calculated from those five pathway modules, matching the modules displayed in Figures 6 and 7. Detailed module definitions and gene membership are provided below.

### Segmentation, cell annotation and anatomical registration

Watershed segmentations were refined using a custom Cellpose2 model trained with manually annotated boundaries; its estimated F1 score was approximately 0.90. The custom model was trained through the Vizgen Postprocessing Tool (VPT) and was selected after comparison with the original watershed segmentation and the default Cellpose model on randomly selected image regions. Cell annotations were assigned with MetaMarkers using the adult mouse-brain reference taxonomy of Yao et al. (2023), using the top 50 markers per cell type at the class level ranked by AUROC. DAPI images and spatial metadata were aligned to Allen CCFv3 using QuickNII and refined by nonlinear alignment in VisuAlign. Final VisuAlign outputs were converted to 16-bit atlas-label images using custom Python scripts. Registered atlas-label images were used to assign cells and transcripts to CCF-defined regions. Cells outside labeled regions and cells with 0 detected transcripts across all genes were excluded; cells with an “unassigned” cell-type annotation from MetaMarkers were excluded from analyses requiring cell-type identity. We normalized gene counts within each cell to 1 across all genes to control for differences in transcript detection between cells. No additional expression-based preprocessing or imputation was applied. Sections were retained as section-level observations and linked to their animal of origin. Registration accuracy was assessed against the corpus callosum, olfactory tubercle and striatal landmarks and independently evaluated by two investigators. A region absent from a section was treated as missing. Distance-resolved neighborhoods were calculated from measured cell coordinates within each section; CCF labels stratified the anatomical analyses but did not generate cell-cell adjacency. Normalized values are compositional within the targeted 500-gene panel and were interpreted as relative expression, not absolute transcript abundance. In animal-level analyses, the two sections from the same mouse did not constitute additional biological replicates. Each section was assigned to its young or old age group and classified as an anterior or posterior sampling level before downstream analysis.

### DDR modules and regional expression-coexpression analysis

Genes were grouped into BER, NER, MMR, DSB, TLS and CoreRep modules according to established functional roles; composite DDR scores integrated the pathway modules used in each analysis by module-level mean.

For regional differential expression analysis, we measured the average cell expression for each gene within each animal, region and cell type to produce per-gene animal-level pseudobulk estimates. Within each cell type and region, old and young pseudobulk values were compared using a paired sign-flip permutation test. The observed mean paired difference was evaluated against a two-sided empirical null generated from 10,000 random sign flips of the per-gene differences (random_state = 42), with a +1 continuity correction. Cell-type-specific P values within a region were combined using Fisher’s method (combine_pvalues, method = ’fisher’) to define the Overall regional P value, replacing any directly pooled-cell estimate for the Overall row. Benjamini-Hochberg FDR correction was applied to the Fisher-combined Overall P values as one family of regions within each module; cell-type-specific P values were reported without FDR correction. The Fisher combination summarized evidence across cell types and did not treat cell types as independent biological replicates; individual cells did not define biological n.

For regional differential coexpression analysis, we measured the average cell expression for each gene within each animal, region and cell type to produce per-gene animal-level pseudobulk estimates. For each module, region and cell type, all possible pairs of genes in the module were enumerated, and Pearson correlations across the animal-level pseudobulk values were calculated separately for young and old samples. Correlations for the same gene pair in the two age groups were Fisher z-transformed and compared by paired t-test, with matched gene pairs as the sampling unit. The resulting P value tested the module-level shift in the distribution of gene-pair correlations between ages. For Overall rows, the same analysis was applied to pseudobulk expression calculated across all cells in each region. Benjamini-Hochberg FDR correction was applied across Overall region-level P values within each module; cell-type-specific P values were reported without FDR correction. Only regions meeting the prespecified minimum cell-count threshold (at least 3 cells) and minimum-animal threshold (at least 3 animals per age group) were tested; other regions were excluded rather than imputed. Gene pairs were the sampling unit for the pathway-level distributional comparison and, because pairs share genes, were not interpreted as independent biological replicates or as evidence for individual regulatory edges. A region absent from an animal was treated as missing and no expression or coexpression value was imputed. For both regional expression and coexpression, the plotted statistic was -log10(q) x sign(mean_young - mean_old), using the FDR-corrected q value. Positive scores indicate higher average values in young animals, whereas negative scores indicate higher average values in old animals. These scores report an age contrast, not a ranking of absolute expression or coordination across regions, and are not effect sizes.

### Inter-module coexpression across matched fine regions

Inter-module analysis used 123 matched fine regions with complete values for all six modules as paired units for inference. For each age, overall within-module coexpression means were correlated across regions. Age-associated changes in correlation were tested by 10,000 within-region young-old label swaps, and 95% confidence intervals were obtained from 10,000 paired regional bootstraps. Benjamini-Hochberg FDR correction was applied across the 15 module pairs; no age-difference edge survived this correction. Spearman correlations and residualization within major anatomical compartments were used as sensitivity analyses. The matched region was the permutation and bootstrap unit; neither cells nor gene pairs were treated as additional biological replicates. Pooled regional coexpression can reflect differences in cellular composition and was not interpreted as evidence of within-cell-type regulatory coordination. The analysis used Overall rows from the matched fine-region table; Allen root and the broad Thalamus parent were excluded. Within-age P values were FDR-corrected across the 15 unique module pairs. Regions were not selected on the basis of a within-module age-effect P value.

### Cellular composition

For each sample and region, cell-type composition was calculated as the proportion of all detected cells, including zero proportions when a sampled region lacked a given cell type. Old and young proportions were compared by two-sided Welch’s t-test; P values were FDR corrected across cell types within each region. For broader regions, cell proportions were recomputed and compared using the same t-test. Analyses were performed using fine anatomical regions and fine cell-type annotations, broad anatomical regions and fine cell-type annotations, and broad anatomical regions and aggregated cell classes. A zero proportion was included only when the anatomical region was sampled in that section but the specified cell type was not detected; a region not present in a section was treated as missing, not as a biological zero. Where fold changes were visualized, a pseudocount of 10^-6 percentage points was added before calculating the log2-transformed ratio of mean old to mean young proportions. The resulting proportions measure relative representation among mapped cells and do not estimate absolute cell density. Sections remained nested within animals; these tests therefore describe variation across sampled sections and do not increase the number of biological replicates. Region-cell-type pairs with fewer than three sampled sections in either age group were not tested. Within each region, cell-type-specific tests were treated as one family; q <= 0.05 was used to define statistical significance.

### Spatial metacells

Spatial metacells were generated independently within each section by k-means clustering of cellular coordinates, targeting approximately 100 cells per metacell. Neuronal expression was pseudobulk-averaged across all neuronal cells in each metacell. To calculate a gene-set score for each metacell, all expressed genes were ranked by percentile, with unexpressed genes excluded. The mean percentile rank of genes in the set among all expressed genes was used as the gene-set score. Cell-type proportions were calculated independently among all cells in each metacell. Immune, astrocyte/ependymal and OPC-oligodendroglial populations were analyzed individually and as a combined group. Scores were calculated for six curated DDR modules and approximately 4,000 GO-derived gene sets containing 2-35 genes, as well as all GO-derived gene sets descending from GO: DNA repair. Selected GO programs were also included independently of the 2-35-gene size range. Composite-DDR scores were calculated as the average gene-set score across all curated DDR modules in each metacell. These scores were used to assess variation in local cell-type composition across neuronal gene-expression states. Composite-DDR-ranked metacells were divided into 50 equal-rank groups and visualized against metacell immune, OPC-oligo, and astro-EPN cell composition. For each curated or GO gene set and age group, target-cell proportions in the highest- and lowest-scoring metacell groups were compared using a two-sided Welch’s independent-samples t-test. The effect size was the log2 fold change in mean cell proportion in the highest-scoring group relative to the lowest-scoring group. Each volcano-plot point therefore reports the significance, magnitude and direction of the local compositional difference associated with one gene set. Within each age group and target-cell-population analysis, all tested gene sets were treated as one family and corrected using the Benjamini-Hochberg procedure; q <= 0.05 defined statistical significance. Metacells are local spatial units nested within sections and animals, not additional biological replicates. These tests quantify associations between local neuronal gene-set scores and local cell composition within this cohort. Rank-based scores reflect relative gene-set expression within the targeted panel and reduce dependence on total transcript detection; they do not measure absolute pathway activity.

### Annular spatial neighborhood analysis

Spatial neighborhood composition was quantified with an annulus-based analysis of age- and DNA damage response (DDR) state-associated differences around defined anchor-cell populations. Analyses were performed on sample-level AnnData objects containing normalized gene-expression measurements, spatial coordinates, anatomical registrations, cell-type annotations and DDR module scores. Cells were retained if they had a nonzero summed normalized expression, were registered to an annotated brain region and, for analyses requiring a cell-type identity, had an assigned MetaMarkers-derived annotation. Gene-expression counts for each cell were normalized to a total of 1 using scanpy.pp.normalize_total(). Module scores were calculated with the rank_among_expressed option of the custom GeneSetScorer.compute_gene_list_score() script. Expressed genes within each cell were converted to percentile ranks; genes with zero expression were excluded rather than assigned a zero rank. The module score was the mean percentile rank of the detected genes belonging to that module. Scores were calculated for five pathway-specific DDR modules: base excision repair (BER), double-strand break repair (DSB), mismatch repair (MMR), nucleotide excision repair (NER) and translesion synthesis (TLS). A Composite-DDR score was calculated from the five pathway modules. Anchor cells were assigned to one of four broad cellular classes: neurons, glia, immune cells or vascular cells; the analyses reported in Figures 6 and 7 focused on neuronal and vascular anchors. For each DDR module independently, anchors were classified as DDR-low or DDR-high using the lower and upper tails of the corresponding module-score distribution, defined as scores at or below the 6th percentile and at or above the 94th percentile, respectively. Percentile cutoffs were calculated across all eligible cells for the corresponding module. The rank-based module score reflects relative within-panel expression and is not a direct measurement of DNA-lesion burden or repair capacity.

For every anchor cell, the surrounding cellular neighborhood was partitioned into ten non-overlapping radial annuli spanning 0-150 µm: 0-5, 5-10, 10-20, 20-30, 30-40, 40-50, 50-75, 75-100, 100-125 and 125-150 µm. Annulus widths increased with distance to retain finer resolution near the anchor while limiting computational cost. Spatial connectivity between each anchor and surrounding cells was determined separately for each annulus using LIANA. Within each annulus, neighborhood composition was calculated as the proportion of cells belonging to each metamarker-defined neighbor cell type relative to the total number of cells present in that annulus. Each anchor cell therefore had an annulus-resolved profile of neighboring cell-type proportions across radial distance. Annular distances were calculated from the measured x-y coordinates within each tissue section. CCF labels were used to stratify the resulting profiles by anatomical region and did not generate the spatial relationships.

### Differential neighborhood composition by age and DDR state

Differential neighborhood composition was evaluated separately by age and DDR state. For age comparisons, anchor cells from young sections were pooled and compared with anchor cells pooled from old sections. For DDR-state comparisons, DDR-low anchors from all sections were pooled and compared with DDR-high anchors from all sections within each age group and DDR module. For each anchor-cell class, neighbor-cell type and annulus, group differences in neighborhood proportions were evaluated using two-sided Mann-Whitney U tests. Effect sizes were quantified using Cohen’s d. Anchor cells remained nested within sections and animals. These Mann-Whitney tests compare anchor-level spatial distributions within the sampled tissue; anchor number does not increase the number of biological replicates. Repeated patterns across sections, regions, modules and neighbor classes were treated as supporting consistency within this cohort, not as independent replication.

For age comparisons, Cohen’s d was oriented as Old minus Young. For DDR-state comparisons, Cohen’s d was oriented as DDR-low minus DDR-high. Accordingly, for the within-age DDR-state comparisons used in the radial summary analyses below, positive Cohen’s d values indicate a greater proportion of the specified neighbor population around DDR-low anchors, whereas negative values indicate a greater proportion around DDR-high anchors. Multiple testing was controlled within each analyzed brain region using Bonferroni correction.

Comparisons were performed for the whole brain and for broad anatomical subdivisions, including cerebral hemisphere, cerebral cortex, cerebral nuclei, brain stem, thalamus, hypothalamus, hippocampal formation, hippocampus, cornu ammonis, dentate gyrus and fiber tracts. The summary visualizations used in the main analysis focused on the whole brain (WB), cerebral cortex (CTX), hippocampal formation (HPF), cerebral nuclei (CNU), thalamus (TH) and hypothalamus (HY).

### Radial integration of annulus-resolved DDR-state effects

To summarize age-dependent spatial organization for visualization, the annulus-resolved Cohen’s d profiles from the within-age DDR-low versus DDR-high comparisons were summarized separately for young and old animals. Only the DDR-state Cohen’s d values calculated within each age group were used for this radial integration; the direct Young-versus-Old Cohen’s d values generated by the primary neighborhood-composition pipeline were not used to calculate the radial summary scores.

For a given brain region, DDR module, anchor class and neighbor class, the age-specific radial association score was calculated as an annulus-width-weighted mean of the signed Cohen’s d values across the available radial bins:

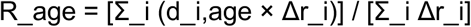

where d_i,age is the DDR-low minus DDR-high Cohen’s d for annulus i in the specified age group and Δr_i is the width of that annulus. The denominator included the widths of annuli with finite Cohen’s d values. Wider annuli therefore received proportionally greater weight than narrower annuli. No Cohen’s d threshold was applied before calculation of the radial association score; all available annuli contributed to the signed summary.

Because Cohen’s d was defined as DDR-low minus DDR-high, positive radial association scores indicate an overall neighborhood bias toward DDR-low anchors across the sampled distances, whereas negative scores indicate an overall bias toward DDR-high anchors. Scores near zero indicate little net directional bias across distance; however, a near-zero summary can also arise when positive and negative annular effects cancel across radial distance and therefore does not necessarily imply absence of spatial structure.

### Aggregation of broad neighbor classes

When a plotted broad neighbor class contained multiple fine-neighbor annotations, an age-specific radial association score was first calculated independently for each fine neighbor annotation and the resulting scores were then averaged using an unweighted arithmetic mean to obtain the broad-class score. The strongest annular effect-size tier and the presence of any qualifying adjusted P value among the contributing fine-neighbor profiles were retained for graphical annotation.

### Age-polarization score

To summarize the age-related change in the radial DDR-state association, an age-polarization score (APS) was calculated for each region, DDR module and anchor-neighbor relationship as the difference between the old and young radial association scores:

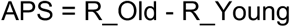

A positive APS therefore indicates that aging shifts the spatial association in the direction of DDR-low anchors, whereas a negative APS indicates a shift in the direction of DDR-high anchors. APS is the change in the integrated effect-size profile between age groups and is distinct from the age-specific radial association scores. Accordingly, identical APS values can arise from different underlying young and old states, including strengthening of an existing association, attenuation of an existing association or reversal across zero.

APS was used as a descriptive summary of the difference between independently estimated age-specific effect-size profiles and was not treated as a formal age-by-DDR-state interaction statistic. Formal inference on such an interaction would require an analysis that retains the appropriate sample- or animal-level replication structure. No independent P value was assigned to APS itself.

### Directional dumbbell and bubble-plot visualization

Directional dumbbell plots were used to display the two age-specific radial association scores for each region and DDR module. Young values were displayed as teal circles and old values as purple squares, with an arrow connecting the young value to the corresponding old value. Positions to the right of zero denote DDR-low-associated neighborhoods and positions to the left of zero denote DDR-high-associated neighborhoods. The arrow shows the young value together with the direction and magnitude of the change to the old value. Arrow direction indicates the transition from the young to the old summary and does not imply a direction of cellular signaling.

Bubble plots provided a compact representation of the same age-associated change. Bubble color encoded the signed APS, with positive values indicating a shift toward DDR-low-associated neighborhoods and negative values indicating a shift toward DDR-high-associated neighborhoods. Bubble area was proportional to the absolute magnitude of APS (|APS|). Bubble plots summarize the age-related change, whereas dumbbell plots display the separate young and old radial association scores.

### Graphical annotation of statistical significance and annular effect size

Statistical significance and annular effect magnitude were displayed as graphical annotations and did not alter the radial association score or APS calculations. For an age-specific point in a directional dumbbell plot, a significance outline was applied when at least one contributing annulus for that age had a Bonferroni-adjusted two-sided Mann-Whitney U-test P value (adj_pval_mw2s_Bonf) of 0.05 or less. Young and old significant points were outlined using age-specific lighter versions of the corresponding marker colors.

Interior stars denoted the maximum absolute annular Cohen’s d observed within the corresponding age-specific radial profile: a small star for maximum |d| of at least 0.20, a medium star for maximum |d| of at least 0.50 and a large star for maximum |d| of at least 0.80. The stars indicate the magnitude of the largest local annular effect and should not be interpreted as significance indicators.

For bubble plots, a significance outline was applied when either age contained at least one contributing annulus with adj_pval_mw2s_Bonf of 0.05 or less. The bubble star reflected the strongest maximum absolute annular Cohen’s d tier observed across the two ages and, where applicable, across contributing fine-neighbor annotations. Significance outlines and effect-size stars report the underlying annulus-level results; bubble position, color and size remain determined by APS.

### Interpretation of radial summaries

The radial association score summarizes the signed annulus-resolved profile across distance but does not retain all spatial detail. Strong effects of opposite sign at proximal and distal radii can partially cancel in the integrated score. Annulus-resolved Cohen’s d profiles therefore provide the most detailed spatial representation; directional dumbbell plots compare the age-specific integrated scores, and APS bubble plots summarize age-related differences across regions and DDR modules.

### Statistical units, nesting and scope of inference

The statistical unit was specified separately for each question. Animal-level pseudobulk values were used for regional gene estimates; matched gene pairs were used for the within-module pathway-level comparison; matched fine regions were used for inter-module age-change inference; section-level proportions were used for the composition analysis; and metacells or anchor-annulus profiles were used for local spatial association. Metacells and anchors are nested spatial observations; their P values were interpreted at the corresponding spatial-unit level within this cohort. Agreement between metacell and annular analyses was interpreted as consistency between analyses of the same cohort, not as independent experimental validation. All DDR-low and DDR-high comparisons are cross-sectional associations; neither spatial proximity, analytical conditioning nor graphical arrows establish causal or signaling direction.

### Software

The annular neighborhood workflow used Scanpy v1.11.5, AnnData v0.11.4, pandas v2.3.3, NumPy v2.2.6, LIANA v1.8.0, Matplotlib v3.10.9, seaborn v0.13.2, plotnine v0.15.7, HNOCA v0.2.1 and Pingouin v0.6.1. Rank-based gene-set scoring, radial association scores, APS values and the associated directional dumbbell and bubble plots were generated with custom Python analysis scripts that will be included with the code release. GeneSetScorer was a custom project script used for rank-based scoring and will be included with the code release; it was not treated as an external software dependency.

### Functional definition of DNA-repair modules

Genes represented in the targeted MERFISH panel were grouped a priori into six functional modules based on established biochemical roles of the encoded proteins, rather than on expression patterns, age effects, spatial distributions or data-driven clustering. Modules were defined at the level of major repair pathways: base excision repair (BER) included factors involved in recognition and repair of small, relatively non-helix-distorting base lesions together with downstream gap processing and ligation; nucleotide excision repair (NER) included factors involved in recognition, unwinding, dual incision and repair synthesis for bulky or helix-distorting lesions; mismatch repair (MMR) included mismatch recognition, excision and repair-synthesis factors; double-strand break repair (DSB) included factors from the major double-strand-break repair pathways, including non-homologous end joining and homologous-recombination/resection functions; translesion synthesis (TLS) included specialized lesion-bypass polymerases and associated factors that permit DNA synthesis across damaged templates; and CoreRep included the core DNA-replication machinery represented in the panel, including clamp/clamp-loader components, replicative polymerases, single-stranded-DNA-binding factors and Okazaki-fragment processing proteins (Krokan and Bjørås, 2013; Marteijn et al., 2014; Jiricny, 2006; Jasin and Rothstein, 2013; Chang et al., 2017; Sale et al., 2012; Burgers and Kunkel, 2017).

### Module membership

**BER:** Ogg1, Nthl1, Neil1, Neil2, Neil3, Ung, Smug1, Mutyh, Mpg, Mbd4, Tdg, Tdg-ps, Apex1, Apex2, Polb, Poll, Fen1, Lig1, Lig3, Xrcc1, Parp1, Parp2, Parp3, Hmgb1, Pcna

**NER:** Xpc, Ddb1, Ddb2, Cul4a, Cul4b, Rbx1 (Roc1), Rad23a, Rad23b, Cetn2, Ercc6, Ercc8, Cdk7, Mnat1, Ccnh, Ercc3, Ercc2, Gtf2h1, Gtf2h2, Gtf2h3, Gtf2h4, Gtf2h5, Ercc5, Ercc4, Ercc1, Xpa, Rpa1, Rpa2, Rpa3

**MMR:** Msh2, Msh3, Msh6, Mlh1, Pms2, Mlh3, Exo1

**DSB:** Xrcc6, Xrcc5, Prkdc, Dclre1c, Polm, Dntt, Lig4, Xrcc4, Nhej1, Rad50, Mre11a, Exo1

**TLS:** Rev1, Polh, Poli, Polk, Poln

**CoreRep:** Pcna, Rfc1, Rfc2, Rfc3, Rfc4, Rfc5, Pold1, Pold2, Pold3, Pold4, Pole, Pole2, Pole3, Pole4, Fen1, Lig1, Rpa1, Rpa2, Rpa3

The initial curation included finer mechanistic subdivisions and cross-pathway roles, but these submodules were not used in the analyses. Because genome-maintenance proteins are frequently multifunctional, module labels were used as functional categories and do not imply exclusive pathway membership. For example, factors such as PCNA, FEN1, LIG1 and RPA participate in replication as well as repair-associated DNA synthesis, EXO1 contributes to both mismatch processing and double-strand-break resection, and CRL4-DDB2 components have NER-linked functions in addition to broader ubiquitin-dependent regulation. When a single module designation was required, the principal pathway assignment was used. CoreRep was treated as a DNA-replication module rather than a canonical DNA-repair pathway. Composite-DDR analyses based on repair pathways therefore used BER, NER, MMR, DSB and TLS, whereas module-coordination analyses included all six modules. In the metacell analysis, the overall DDR score used to rank metacells was the mean of all six curated module scores, as specified in the Spatial metacells subsection; the annular Composite-DDR score was the mean of BER, NER, MMR, DSB and TLS.

